# Genomically mined acoustic reporter genes enable real-time *in vivo* monitoring of tumors and tumor-homing probiotics

**DOI:** 10.1101/2021.04.26.441537

**Authors:** Robert C. Hurt, Marjorie T. Buss, Mengtong Duan, Katie Wong, Mei Yi You, Daniel P. Sawyer, Margaret B. Swift, Przemysław Dutka, Pierina Barturen-Larrea, David R. Mittelstein, Zhiyang Jin, Mohamad H. Abedi, Arash Farhadi, Ramya Deshpande, Mikhail G. Shapiro

## Abstract

A major outstanding challenge in the fields of biological research, synthetic biology and cell-based medicine is visualizing the function of natural and engineered cells noninvasively inside opaque organisms. Ultrasound imaging has the potential to address this challenge as a widely available technique with a tissue penetration of several centimeters and spatial resolution below 100 μm. Recently, the first genetically encoded acoustic reporters were developed based on bacterial gas vesicles to link ultrasound signals to molecular and cellular function. However, the properties of these first-generation acoustic reporter genes (ARGs) resulted in limited sensitivity and specificity for imaging gene expression *in vivo.* Here, we describe second-generation ARGs with greatly improved acoustic properties and expression characteristics, identified through a phylogenetic screen of candidate gene clusters from diverse bacteria and archaea. The resulting constructs offer major qualitative and quantitative improvements, including much stronger ultrasound contrast, the ability to produce nonlinear signals distinguishable from background tissue, and stable long-term expression. We demonstrate the capabilities of these next-generation ARGs by imaging *in situ* gene expression in mouse models of breast cancer and tumor-homing therapeutic bacteria, noninvasively revealing the unique spatial distributions of tumor growth and colonization by therapeutic cells in living subjects and providing real-time guidance for interventions such as needle biopsies.

## INTRODUCTION

Basic biological research, *in vivo* synthetic biology and the development of cell-based medicine require methods to visualize the functions of specific cells deep inside intact organisms. In this context, widely used optical techniques based on fluorescent and luminescent proteins have limited utility due to the scattering and absorption of light by tissue.^1^ In contrast, ultrasound is a widely used technique for deep-tissue imaging, providing sub-100 μm spatial resolution and penetrating several cm into tissue.^2^ The relative simplicity and low cost of ultrasound make it widely accessible for both research and clinical medicine, while recently developed super-resolution methods^3,4^ push its spatial resolution below 10 μm. Recently, the first genetically encodable reporters for ultrasound^5–7^ were introduced based on gas vesicles (GVs): air-filled protein nanostructures encoded by clusters of 8-20+ genes, which evolved as flotation devices in a wide range of mostly aquatic bacteria and archaea.^8,9^ The low density and high compressibility of their air-filled interiors compared to surrounding tissues allow GVs to scatter sound waves and thereby produce ultrasound contrast when heterologously expressed as acoustic reporter genes (ARGs) in genetically engineered bacteria^6^ or mammalian cells.^7^

Despite the conceptual promise of first-generation ARGs, several limitations prevent their widespread use for monitoring bacterial or mammalian gene expression *in vivo*. First-generation bacterial ARGs^6^ cannot scatter ultrasound nonlinearly (making them difficult to distinguish from background tissues), express poorly at 37°C, and are too metabolically burdensome for *in situ* expression *in vivo*. Likewise, first-generation mammalian ARGs^7^ also produced only linear contrast, and cell-to-cell variability in their expression and burden meant that they could only be imaged robustly with ultrasound in clonally selected cell lines stimulated with potent epigenetic reagents. In both cases, the lack of nonlinear signal had to be circumvented by destructive ultrasound pulse sequences, which destroyed the GVs and limited dynamic imaging.^10^ Just as the widespread use of fluorescent proteins did not take off until the development of enhanced versions of GFP, an analogous breakthrough is required for acoustic proteins to become widely useful in *in vivo* biological research and potential clinical applications.

Motivated by this challenge, we sought nextgeneration ARGs that, when expressed heterologously in either probiotic bacterial strains or mammalian cancer cell lines, could produce GVs with strong nonlinear ultrasound contrast and enable robust, sustained expression under physiological conditions. These qualities would enable longterm noninvasive imaging of gene expression in a broad range of *in vivo* applications. We hypothesized that a genomic mining approach – previously applied to improving fluorescent proteins^11–14^, opsins^15–17^ Cas proteins^18–22^, and other biotechnology tools^23–28^ would yield ARGs with improved properties, which could be further optimized through genetic engineering. By cloning and screening 15 distinct polycistronic operons from a diverse set of GV-expressing species representing a broad phylogeny, we identified two GV gene clusters – from *Serratia sp.* 39006 and *Anabaena flos-aquae* – that produce vastly more linear and nonlinear acoustic contrast than previously tested clusters when expressed in several types of bacteria and mammalian cells, respectively. The bacterial ARG adapted from *Serratia* sp. 39006 (bARG_Ser_), when expressed in the widely used probiotic bacterium *E. coli* Nissle 1917 (EcN), enabled noninvasive ultrasound imaging of these probiotic agents colonizing tumors, providing direct visualization of the microscale *in vivo* distribution of this rapidly emerging class of anti-cancer therapy. The mammalian ARG adapted from *A. flos-aquae* (mARG_Ana_), when expressed in human breast cancer cells, enabled both the noninvasive, *in situ* microscale imaging and long-term monitoring of heterologous gene expression in developing orthotopic tumors, and the ultrasound-guided biopsy of a genetically defined subpopulation of these tumor cells. The properties and performance of these second-generation ARGs represent a fundamental advance in the utility of acoustic proteins for a wide range of *in vivo* research.

## RESULTS

### Genomic mining of gas vesicle gene clusters reveals homologs with improved ultrasound performance in E. coli

GVs are encoded by polycistronic gene clusters comprising one or more copies of the primary structural gene *gvpA* and 7 to 20+ other genes encoding minor constituents, assembly factors or reinforcing proteins, which together help assemble the GVs’ protein shells.^9^ Hundreds of organisms have GV genes in their genomes, but only a small subset have been shown to produce GVs. Given the labor involved in cloning and testing polycistronic clusters, we limited our phylogenetic search to organisms with confirmed GV production and sequenced operons (**Table S1**). We selected 11 representative species, broadly sampling phylogenetic space, cluster architecture and organismal characteristics (*i.e*., halophilic, thermophilic and mesophilic) (**Fig. 1a** and **Fig. S1**). We obtained each species from culture repositories, amplified GV operons from their genomes, and cloned them into a bacterial expression vector.

**Figure 1.**
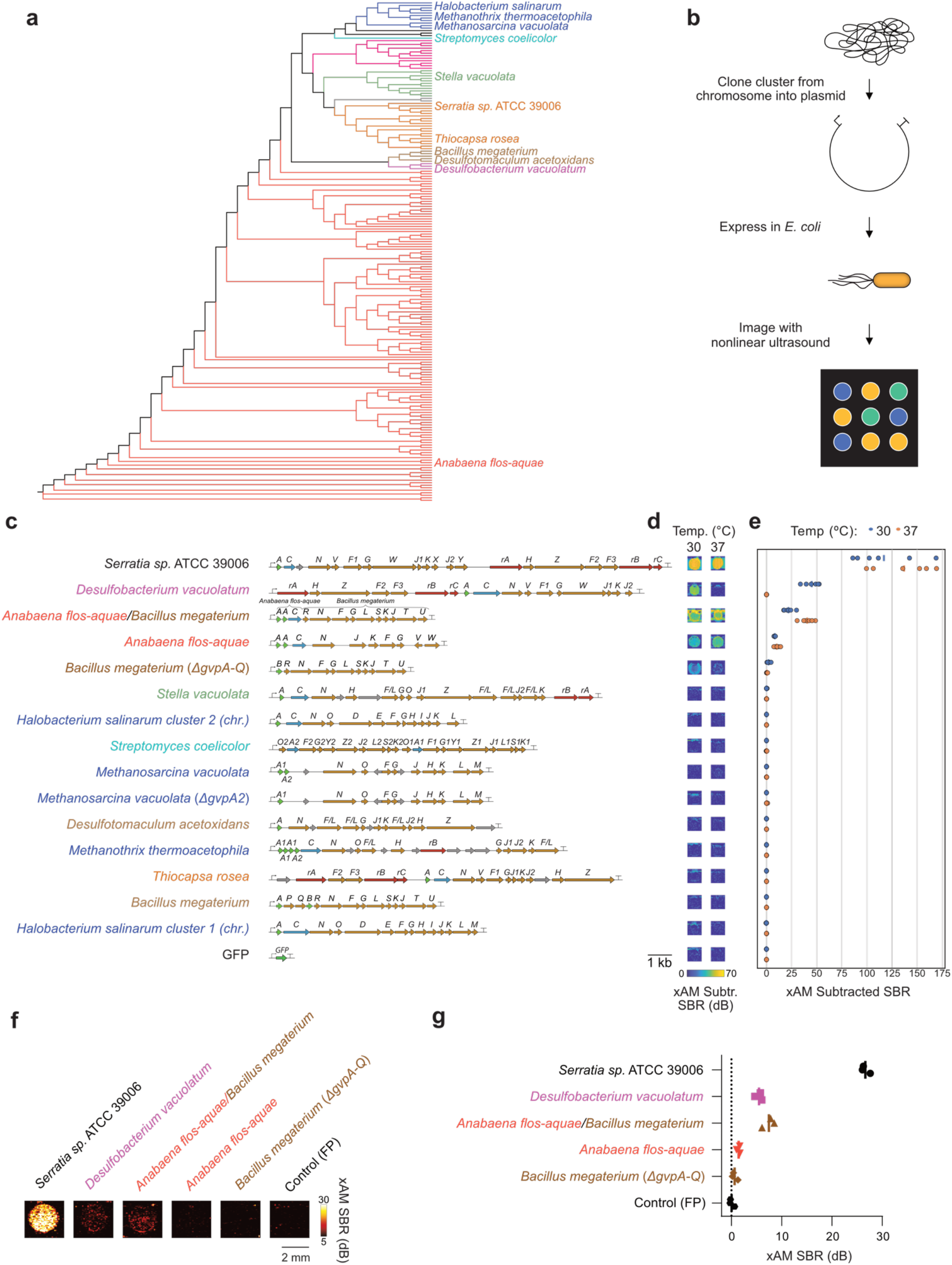
Genomic mining of gas vesicle gene clusters reveals homologs with nonlinear ultrasound contrast in *E. coli.* (**a**) 16S phylogenic tree of known GV-producing organisms, with the species from which GV genes were cloned and tested in this study indicated by name. See **Fig. S1** for the fully annotated phylogenic tree. *B. megaterium* and *S’. coelicolor* were not reported to produce GVs, but we tested their GV gene clusters here based on previous experiments in *E. coli*^3^ and to broadly sample the phylogenetic space. (**b**) Workflow for testing GV clusters. Selected GV gene clusters were expressed in BL21 (DE3) *E. coli* by growing patches of cells on plates containing the inducer IPTG, and the patches were then imaged with nonlinear ultrasound (xAM). (**c-e**) Diagrams of the GV gene clusters tested in *E. coli* (c), differential xAM images of representative patches (d), and quantification of the differential xAM signal-to-background ratio (SBR) of the patches (n=6 biological replicates) (e). (**f-g**) Representative xAM images (f) and quantification of the xAM SBR (n=3 biological replicates, each with 2 technical replicates; lines represent the mean) (g) for the top 5 GV-producing clusters expressed in *E. coli* at 30’>C on solid media and normalized to 5 × 10^9^ cells/mL in agarose phantoms, imaged at 1.74 MPa. See **Fig. S7a-b** for the ultrasound signal at varying acoustic pressures and **Fig. S7c** for the corresponding BURST data.

We then expressed each operon in confluent *E. coli* patches at several temperatures and inducer concentrations (**Fig. 1b**), comparing them to two bacterial ARG constructs previously shown to work in *E. coli*^6^ – bARG1 *(Anabaena flos-aquae/Bacillus megaterium* hybrid) and *Bacillus megaterium ΔgvpA-Q* as well as the full *Bacillus megaterium* gene cluster (**Fig. 1c-g**, **Fig. S2a-e**, **Fig. S3a-c**, **Fig. S4-6**). We scanned these patches using a home-built robotic ultrasound imaging apparatus, applying a cross-propagating amplitude modulation pulse sequence (xAM).^29^ This pulse sequence enhances signals specific to nonlinear contrast agents such as GVs while cancelling linear background scattering. Importantly, unlike pulse sequences that relied on the irreversible collapse of GVs to obtain GV-specific contrast^6,7,10^, xAM is nondestructive. In addition, we examined the optical opacity of the patches, which can be increased by sufficient levels of GV expression. Of the 15 gene clusters tested, only 3 showed significant xAM signal when expressed at 37°C, and 5 showed significant xAM signal at 30°C (**Fig. 1, c-e**). Even though all operons tested are from organisms reported to produce GVs in their native hosts, only the *A. flos-aquae, B. megaterium ΔgvpA-Q* bARG1, *Desulfobacterium vacuolatum,* and *Serratia sp.* 39006 *(Serratia)* clusters produced detectable GVs heterologously in *E. coli*. Several other operons produced a small amount of ultrasound contrast under certain conditions, which did not arise from GV expression but reflected a rough patch morphology likely due to cellular toxicity (**Fig. S3d)**. The failure of most tested gene clusters to produce GVs in *E. coli* is not surprising given the complexity of polycistronic heterologous expression, which requires each component to fold and function properly in a new host with a potentially different cytoplasmic environment, growth temperature and turgor pressure.^30,31^ In addition, it is possible that some genes included in the clusters act as cis-regulators,^9,30,32–34^ limiting expression absent a specific trans input, or that some additional genes are required beyond the annotated operons.

In patch format, the strongest acoustic performance was observed with the genes from *Serratia*, bARG1, *A. flos-aquae*, *B. megateriaum,* and *D. vacuolatum.* Because patch experiments do not control for the density of cells in each sample, we further compared the performance of these clusters in resuspended samples. Each operon was expressed on solid media at 30°C – a temperature at which all five operons produced GVs – then scraped, resuspended, and normalized for cell concentration. These samples were imaged in hydrogels using both xAM (**Fig. 1, f-g** and **Fig. S7, a-b**) and a more sensitive but destructive imaging method called BURST^35^ (**Fig. S7c**), and examined optically with phase-contrast microscopy (PCM), which reveals the presence of GVs due to the refractive index difference between GVs and water^36,37^ (**Fig. S7d**). Three of the clusters produced xAM signals, and all clusters produced BURST signals significantly stronger than the negative control. All clusters except *A. flos-aquae* exhibited sufficient GV expression to be visible by PCM.

Cells expressing the *Serratia* cluster produced the strongest ultrasound signals, 19.2 dB above the next brightest cluster, bARG1, under xAM imaging at an applied acoustic pressure of 1.74 MPa: an 83-fold gain in signal intensity (**Fig. 1f**). Additionally, PCM images (**Fig. S7d**) showed that cells expressing the *Serratia* cluster had the highest levels of GV expression, as also seen in whole-cell transmission electron microscopy (TEM) (**Fig. S7e**). Based on the large improvement in ultrasound contrast provided by the *Serratia* GV operon relative to the other gene clusters, we selected this operon for further optimization as a second-generation bacterial ARG.

Because overexpression of any protein imposes a metabolic demand on the host cell,^38–40^ we reasoned that deletion of non-essential genes could improve GV expression from the *Serratia* cluster, and therefore the xAM signal. Previous work showed that deletions of *gvpC, gvpW, gvpX, gvpY, gvpH,* or *gvpZ* preserve GV formation in the native organism.^30^ We tested these deletions, as well as the deletion of an unannotated hypothetical protein (Ser39006_001280) encoded between the *gvpC* and *gvpN* coding sequences (**Fig. S8a**). When expressed in *E. coli,* deletions of *gvpC, gvpH, gvpW, gvpY,* or *gvpZ* reduced or eliminated xAM signal (**Fig. S8, b-c**) and patch opacity (**Fig. S8d**). Deletion of *gvpX* increased xAM signal but decreased opacity, and deletion of Ser39006_001280 increased both xAM signal and opacity. Based on these results, we selected the *Serratia* ΔSer39006_001280 operon for subsequent *in vitro* and *in vivo* experiments. We call this new genetic construct bARG_Ser_ a bacterial acoustic reporter gene derived from *Serratia*.

### bARG_Ser_ shows robust expression, contrast, and stability in E. coli Nissle

We transferred bARG_Ser_ into EcN, a strain of *E. coli* that is widely used in *in vivo* biotechnology applications due to its ability to persist in the gastrointestinal tract and colonize tumors.^41–43^ EcN has been intensely investigated as a chassis for anti-tumor therapy delivery, and is currently the most commonly used species in this application.^44–47^ We first tested three different inducible promoter architectures in EcN on solid media at 37°C, examining xAM ultrasound contrast and patch opacity as a function of inducer concentration. We found that the L-arabinose-inducible pBAD promoter provided the most robust control over GV expression without obvious burden (**Fig. 2, a-b, Fig. S9**). Based on these results, we selected the pBAD-bARG_Ser_ EcN strain for subsequent experiments.

**Figure 2.**
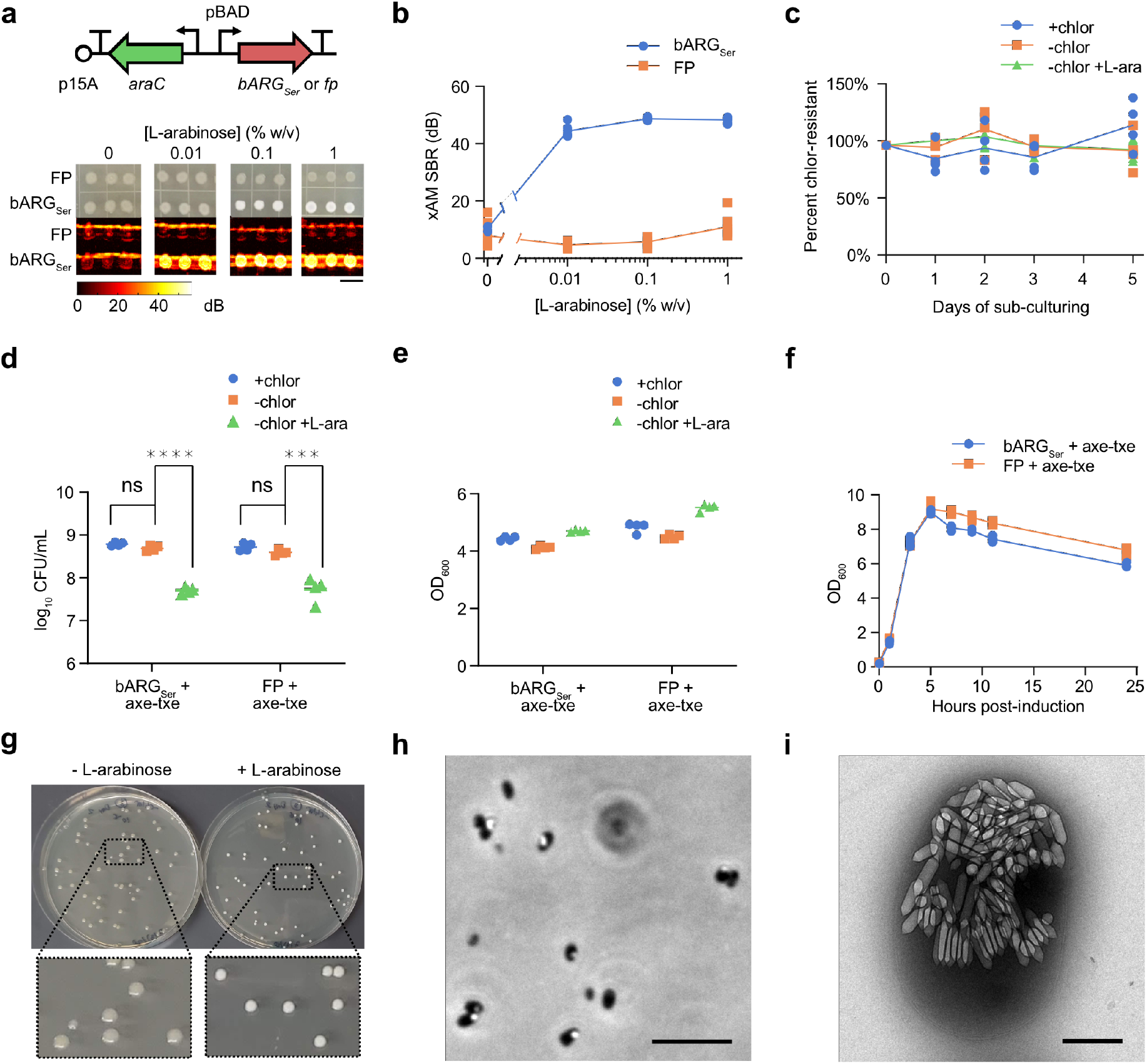
Genetic engineering and expression of bARG_Ser_ in the probiotic bacterium *E. coli* Nissle (EcN). (**a**) Diagram of the arabinose-inducible construct used to express bARG_Ser_ in EcN (top), and optical and xAM images of bARG_Ser_-expressing or FP-expressing patches of EcN on solid media with varying L-arabinose concentrations at 37°C (bottom). The scale bar is 1 cm. See **Fig. S9** for the corresponding results with IPTG-inducible and aTc-inducible constructs. (**b**) Quantification of the xAM SBR of all patches from the experiment in (a) versus the L-arabinose concentration (n=8 biological replicates). (**c**) Diagram of the construct from (a) with the toxin-antitoxin stability cassette Axe-Txe^48^ added to enable plasmid maintenance in the absence of antibiotics (top), and verification of plasmid maintenance *in vitro* (bottom). The percentage of chloramphenicol-resistant colonies was measured during daily sub-culturing into LB media with 25 μg/mL chloramphenicol (+chlor), without chloramphenicol (−chlor), or without chloramphenicol and with 0.1% (w/v) L-arabinose (−chlor +L-ara) using pBAD-bARG_Ser_-AxeTxe EcN (n=4 biological replicates). The percentage of chloramphenicol-resistant colonies was calculated by dividing the number of colonies on plates with chloramphenicol by the number of colonies on plates without chloramphenicol. (**d-e**) Colony forming units (CFUs) per mL of culture on chloramphenicol plates (d) and optical density at 600 nm (e) of pBAD-bARG_Ser_-AxeTxe EcN and pBAD-FP-AxeTxe EcN cultures 24 hours after sub-culturing into LB media with the same conditions as in (c). Asterisks indicate statistical significance by two-tailed, unpaired Student’s t-tests (**** = p < 0.0001, *** = p < 0.001, ns = no significance); n=4 biological replicates. (**f**) OD_600_ versus time after inducing pBAD-bARG_Ser_-AxeTxe (bARG_Ser_ + axe-txe) and pBAD-FP-AxeTxe (FP + axe-txe) EcN strains with 0.1% (w/v) L-arabinose in liquid culture at 37°C (n=4 biological replicates). Between 5 and 24 hours post-induction, when the OD_600_ of all cultures decreased, the OD_600_ of FP-expressing cultures was slightly higher than that of the bARG_Ser_-expressing cultures, likely due to expression of red fluorescent protein which is known to absorb light at 600 nm.^83^ (**g**) Representative image of colonies from the experiment in (c) on chloramphenicol plates with (right) and without (left) 0.1% (w/v) L-arabinose. The opacity of the colonies on plates with L-arabinose indicates bARG_Ser_ expression and was used to screen for mutants deficient in bARG_Ser_ expression (see **Fig. S11**). (**h-i**) Representative phase contrast microscopy (h) and transmission electron microscopy (i) images of pBAD-bARG_Ser_-AxeTxe EcN cells grown on plates with 0.1% (w/v) L-arabinose at 37°C. Scale bars are 10 μm (h) and 500 nm (i). Curves and lines represent the mean for b-f.

To ensure that the pBAD-bARG_Ser_ plasmid is maintained in the absence of antibiotic selection, as required in certain *in vivo* applications, we added the toxin-antitoxin stability cassette Axe-Txe.^48^ This enabled the pBAD-bARG_Ser_-AxeTxe plasmid to be maintained in EcN for up to 5 days of daily sub-culturing in liquid media without antibiotics, both with and without induction of ARG expression (**Fig. 2c**).

The expression of most heterologous genes, including widely used reporter genes such as fluorescent proteins (FPs), results in some degree of metabolic burden on engineered cells.^39,40,49^ Consistent with this expectation, the induction of pBAD-bARG_Ser_ EcN resulted in reduced colony formation to an extent similar to the expression of a FP (**Fig. 2d** and Fig. S10a), even as the culture density measured by OD_600_ remained relatively unchanged (**Fig. 2e** and **Fig. S10b**). When the OD_600_ was measured after inducing cultures with L-arabinose, the growth curves of bARG_Ser_-expressing and FP-expressing EcN were indistinguishable during the growth phase (0 to 5 hours), indicating that the two strains have similar growth rates (**Fig. 2f**). Collectively, these results suggest that overexpression of bARG_Ser_ using the pBAD expression system in EcN is not significantly more burdensome than that of FPs, which are widely accepted as relatively non-perturbative indicators of cellular function.

To further examine the genetic stability of bARG_Ser_ constructs, we plated cells from daily sub-cultures onto agar with 0.1% (w/v) L-arabinose and examined colony opacity (**Fig. 2g**) as a measure of retained GV expression. Of a total of 3824 colonies, nearly all were opaque (**Fig. 2g**), with GV expression confirmed by PCM and TEM (**Fig. 2, h-i**). Only 11 colonies (<0.3% after ~35 cell generations) exhibited a reduced opacity (**Fig. S11a**), representing a mutated phenotype confirmed by growing these cells on fresh media (**Fig. S11b**). PCM revealed that these rare mutants still produced GVs, but at lower levels than non-mutants. These results indicate that mutational inactivation of GV production is not a major issue for pBAD-bARG_Ser_-AxeTxe EcN under typical conditions.

After establishing construct stability, we characterized the acoustic properties of bARG_Ser_-expressing EcN. For cells induced in liquid culture with 0.1% L-arabinose for 24 hours and suspended at 10^9^ cells/mL in agarose phantoms, an xAM signal was detected at acoustic pressures above 0.72 MPa, rising with increasing pressure up to the tested maximum of 1.74 MPa (**Fig. 3a**). To characterize the physical stability of GVs in these cells during ultrasound exposure, we measured the xAM signal over time at a series of increasing acoustic pressures (**Fig. 3b** and **Fig. S12**). The xAM signal was steady at pressures up to 0.96 MPa, above which we observed a slow decrease, indicating that some of the GVs gradually collapsed despite sustained high xAM signals. We also imaged the cells with parabolic pulses, which can transmit higher pressures than xAM, and thus can be helpful *in vivo* to compensate for attenuation at tissue interfaces. When imaged with parabolic B-mode at varying acoustic pressures, the GVs started to collapse slowly at 1.02 MPa and more rapidly at 1.33 MPa and above (**Fig. 3c**). Based on these results, an acoustic pressure of 1.29 MPa was selected for xAM and 1.02 MPa for parabolic AM (pAM) imaging in subsequent experiments to obtain the strongest signals while minimizing GV collapse.

**Figure 3.**
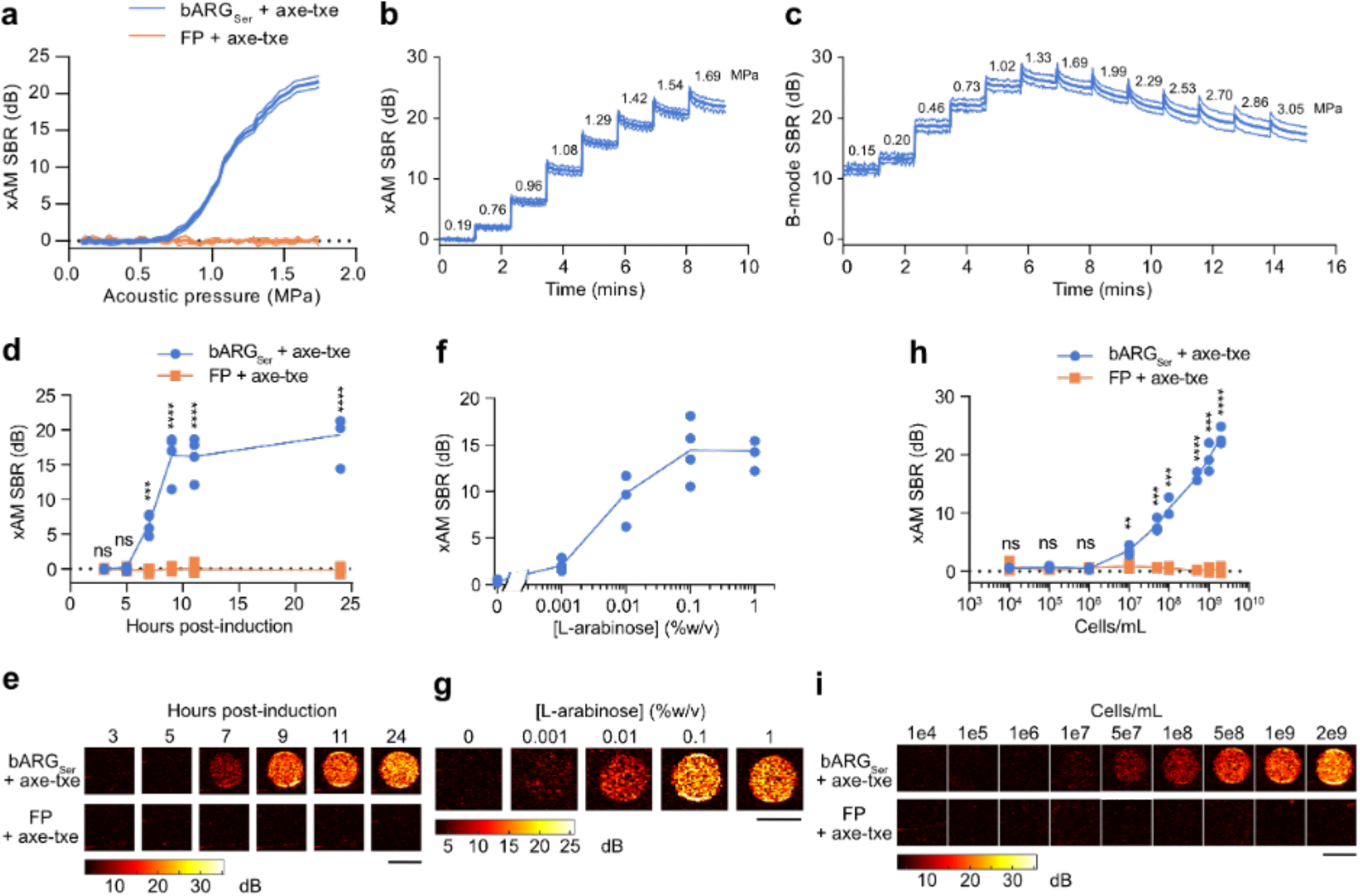
Acoustic characterization of bARG_Ser_-expressing EcN *in vitro*. (**a**) xAM SBR as a function of transmitted acoustic pressure for bARG_Ser_-expressing and FP-expressing EcN. (**b-c**) xAM (b) and parabolic B-mode (c) SBRs measured over time when the transmitted acoustic pressure was increased approximately every 70 sec as indicated by the numbers above the curve for bARG_Ser_-expressing EcN. Ultrasound was applied at a pulse repetition rate of 86.8 Hz. For (a-c), pBAD-bARG_Ser_-AxeTxe EcN were induced with 0.1% (w/v) L-arabinose for 24 hours at 37°C in liquid culture, and were then normalized to 10^9^ cells/mL in agarose phantoms for ultrasound imaging. Bold lines represent the mean and thin lines represent ± standard deviation; n=3 biological replicates, each with 2 technical replicates. (**d-e**) xAM ultrasound SBR (d) and corresponding representative ultrasound images (e) at several time points after inducing pBAD-bARG_Ser_-AxeTxe (bARG_Ser_ + axe-txe) and pBAD-FP-AxeTxe (FP + axe-txe) EcN strains with 0.1% (w/v) L-arabinose in liquid culture at 37°C. (**f-g**) xAM ultrasound SBR (f) and corresponding representative ultrasound images (g) using varying L-arabinose concentrations to induce pBAD-bARG_Ser_-AxeTxe EcN in liquid culture at 37°C for 24 hours. (**h-i**) xAM ultrasound SBR (h) and corresponding representative ultrasound images (i) of varying concentrations of pBAD-bARG_Ser_-AxeTxe or pBAD-FP-AxeTxe EcN cells induced for 24 hours at 37°C with 0.1% (w/v) L-arabinose in liquid culture. For (e, g, i) scale bars are 2 mm. For (d-g), cells were normalized to 10^9^ cells/mL in agarose phantoms for ultrasound imaging. For (d, f, h), each point is a biological replicate (n=4 for d and f; n=3 for h) that is the average of at least 2 technical replicates, and curves represent the mean. Asterisks represent statistical significance by two-tailed, unpaired Student’s t-tests (**** = p<0.0001, *** = p<0.001, ** = p<0.01, ns = no significance).

Next, to characterize the dynamics and inducibility of bARG_Ser_ in EcN and determine the ultrasound detection limit of bARG_Ser_-expressing EcN, we measured xAM signal as a function of induction time, inducer concentration, and cell concentration. At a density of 10^9^ cells/mL, xAM signal could first be observed 7 hours after induction with 0.1 % L-arabinose and leveled off by 9 hours post-induction (**Fig. 3, d-e**). Keeping the induction time constant at 24 hours while varying the L-arabinose concentration, GV expression was detected with as little as 0.001% L-arabinose, and the highest ultrasound signal was observed for 0.1-1% L-arabinose (**Fig. 3, f-g**). When cells induced for 24 hours with 0.1% L-arabinose were diluted, they were detectable by ultrasound down to 10^7^ cells/mL (**Fig. 3, h-i**). Critically, this detection was achieved non-destructively with nonlinear imaging, compared to previous bacterial ARGs, which required a destructive linear imaging approach.^6^ The bARG_Ser_ xAM signal was proportional to the cell concentration between 10^7^ cells/mL and 2 × 10^9^ cells/mL (**Fig. 3, h-i**). We also imaged the cells using BURST imaging, which provides greater sensitivity at the cost of collapsing the GVs.^35^ BURST enabled bARG_Ser_-expressing EcN to be detected as early as 3 hours post-induction (**Fig. S13, a-b**), with as little as 0.001% L-arabinose (**Fig. S13, c-d**), and at a density as low as 10^5^ cells/mL (**Fig. S13, e-f**). Taken together, our *in vitro* experiments indicated that the reporter gene construct pBAD-bARG_Ser_-AxeTxe is robust and stable in EcN and enables gene expression in these cells to be imaged with high contrast and sensitivity.

To test whether this ARG construct can function in other bacterial species, we transformed pBAD-bARG_Ser_-AxeTxe into an attenuated strain of *Salmonella enterica* serovar Typhimurium, which has been used in bacterial anti-tumor therapies.^50^,^51^ After inducing expression with L-arabinose, *S. Typhimurium* cells produced GVs, as observed with PCM (**Fig. S14a**), and generated strong contrast in xAM imaging (**Fig. S14b-c**). *S. Typhimurium* cells exhibited similar detection limits to EcN with both xAM and BURST imaging (**Fig. S14d-k**).

### bARG_Ser_ enables in situ imaging of tumor-colonizing bacteria

Tumor-homing bacteria are a major emerging class of cancer therapy, taking advantage of the ability of cells such as EcN to infiltrate tumors and proliferate in their immunosuppressed microenvironment. Major synthetic biology efforts have been undertaken to turn tumor-homing EcN cells into effective therapies for solid tumors.^44–47^ However, despite the promise of this technology and the importance of appropriate microscale biodistribution of the bacteria inside tumors, no effective methods currently exist to visualize this biodistribution *in situ* in living animals.

To test the ability of bARG_Ser_ to overcome this limitation, we formed subcutaneous MC26 tumors in mice and, when the tumors reached a substantial size, intravenously injected EcN cells containing the pBAD-bARG_Ser_-AxeTxe plasmid, giving the bacteria 3 days to home to and colonize the tumors. We then induced GV expression and imaged the tumors with ultrasound (Fig. 4a). In all tumors colonized by bARG_Ser_-expressing EcN, we observed pAM, BURST, and xAM ultrasound contrast one day after induction with L-arabinose (**Fig. 4b** and **Fig. S15a**). The signals were localized to the core of the tumor and concentrated at the interface between live and necrotic tissue, where the EcN primarily colonized, as confirmed with subsequent tissue histology (**Fig. 4f-i** and **Fig. S16**). This biodistribution reflects the immune-privileged environment of the necrotic tumor core and has been observed previously only with post-mortem *ex vivo* optical methods.^42,52^

**Figure 4.**
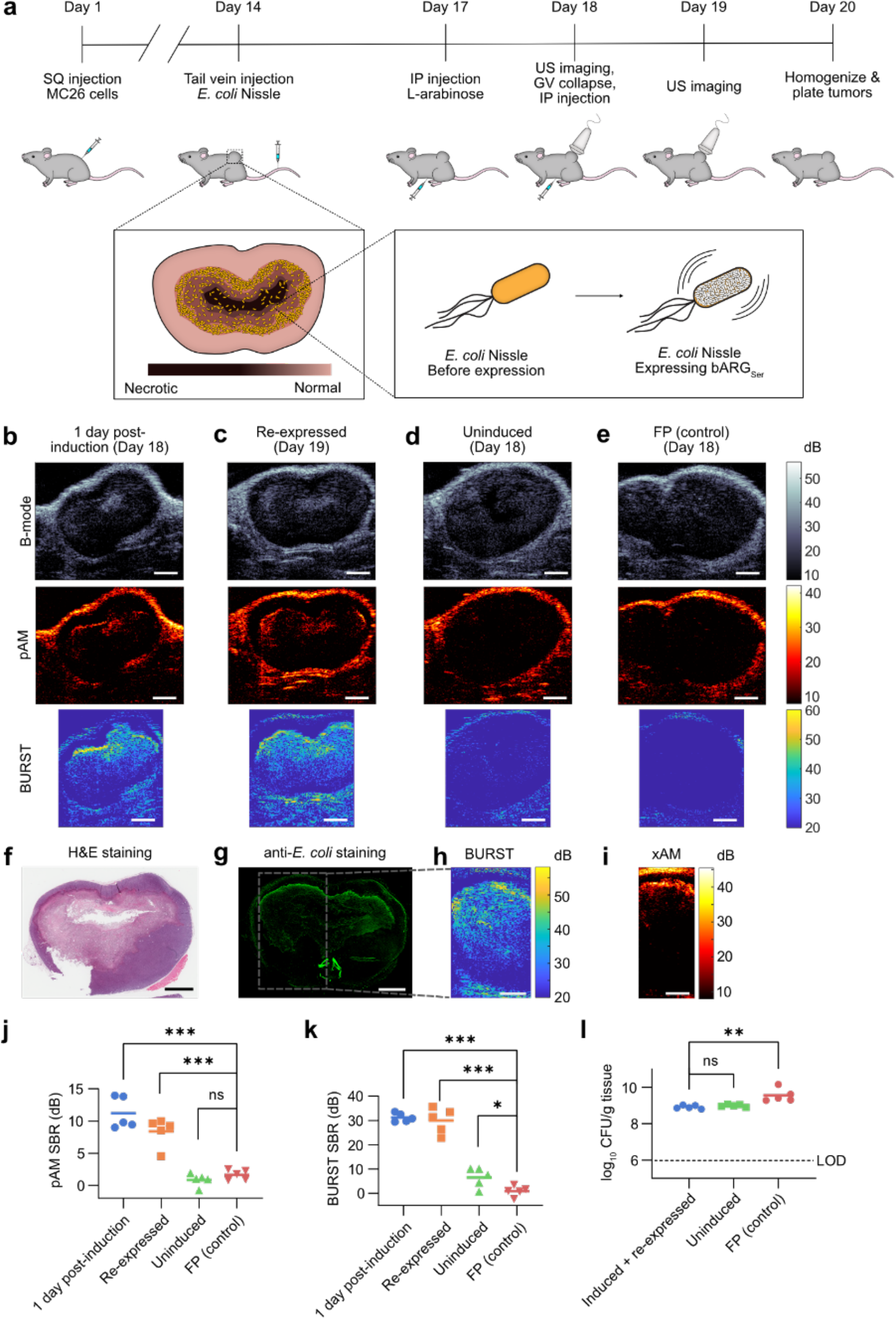
*In situ* bARG_Ser_ expression enables ultrasound imaging of tumor colonization by EcN. (**a**) Diagram of the *in vivo* protocol for assessing *in situ* bARG_Ser_ expression in tumors. Mice were injected subcutaneously (SQ) with MC26 cancer cells on day 1 and the tumors were allowed to grow for 14 days. Mice were then injected with *E. coli* Nissle (EcN) carrying either pBAD-bARG_Ser_-AxeTxe or pBAD-FP-AxeTxe plasmids via the tail vein. After allowing 3 days for the EcN to colonize the tumors, bARG_Ser_ or FP expression was induced by injecting L-arabinose intraperitoneally (IP) on day 17. The next day, at least 24 hours after induction, tumors were imaged with ultrasound. Subsequently, all the GVs in the tumors were collapsed by applying maximum acoustic pressure (3.0 MPa) throughout the tumor. L-arabinose was then injected again to re-induce bARG_Ser_ expression, and tumors were again imaged with ultrasound at least 24 hours later. The next day (day 20), all mice were sacrificed, and their tumors were homogenized and plated on selective media to quantify the levels of EcN colonization. In separate experiments for histological analysis, mice were sacrificed on day 18 directly after ultrasound imaging. (**b-d**) Representative B-mode, parabolic AM (pAM), and BURST ultrasound images of tumors colonized by pBAD-bARG_Ser_-AxeTxe EcN at least 24 hours after induction with L-arabinose on day 18 (b), at least 24 hours after collapse and re-induction (day 19) (c), or uninduced on day 18 (d). (**e**) Representative B-mode, pAM, and BURST ultrasound images of tumors colonized by pBAD-FP-AxeTxe EcN at least 24 hours after induction with L-arabinose on day 18. (**f-g**) Optical images of tissue sections stained with H&E (f) or anti-*E. coli* antibodies (g) from a tumor colonized by bARG_Ser_-expresssing EcN after ultrasound imaging on day 19. (**h-i**) BURST (h) and xAM (i) ultrasound images of the same tumor as in (f-g), with the boxed region showing the approximate BURST imaging region in the tissue section. Scale bars in (b-i) represent 2 mm. (**j-k**) Quantification of the pAM (j) and BURST (k) SBR for the same conditions in (b-e). (**l**) Colony forming units (CFUs) per gram of tissue from tumors homogenized and plated on day 20. For j-l, points represent each mouse (n=5), and lines represent the mean of each group. The dotted line indicates the limit of detection (LOD). Asterisks represent statistical significance by two-tailed, unpaired Student’s t-tests (*** = p<0.001, ** = p<0.01, * = p<0.05, ns = no significance). See **Fig. S13** for representative xAM ultrasound images for the conditions in (b-d), and **Fig. S14** for more histological images of tissue sections from tumors colonized with pBAD-bARG_Ser_-AxeTxe EcN.

Furthermore, after applying 3 MPa of acoustic pressure throughout the tumor to collapse all the GVs, reinjecting the mice with L-arabinose inducer, and allowing ≥24 hours for re-expression, similar ultrasound signals were observed in all tumors colonized by bARG_Ser_-expressing EcN (**Fig. 4c** and **Fig. S15b**). This result shows that bARG_Ser_ can be used to visualize dynamic gene expression at multiple timepoints. Absent L-arabinose induction, no xAM or pAM ultrasound signals were observed from bARG_Ser_-containing EcN (**Fig. 4d** and **Fig. S15c**); likewise, no xAM or pAM ultrasound signals were seen in tumors colonized by FP-expressing EcN (**Fig. 4e,j** and **Fig. S15, d-e**). Low levels of BURST signal were observed in uninduced animals (**Fig. 4k**), likely due to small amounts of L-arabinose present in the diet combined with BURST imaging’s high sensitivity.

To quantify tumor colonization, at the end of the experiment (day 20 in **Fig. 4a**) all mice were euthanized and their tumors were homogenized and plated on selective media. Tumors from all groups of mice (n=5 for induced and reexpressed bARG_Ser_, n=5 for uninduced bARG_Ser_, and n=5 for FP) contained more than 7 × 10^8^ CFU/g tissue (**Fig. 4l**), indicating that the EcN can persist at high levels in tumors for at least 6 days after IV injection regardless of bARG_Ser_ expression, collapse, and re-expression. The somewhat higher density of FP-expressing EcN suggested that maintenance of the smaller pBAD-FP-AxeTxe plasmid (7.2 kb versus 23.2 kb for pBAD-bARG_Ser_-AxeTxe), which would impose less burden on the cell,^53–56^ may be easier in this *in vivo* context where oxygen and other nutrients are limited. Negligible mutational silencing was observed in EcN plated from tumor samples (**Fig. S17, a-c**).

Taken together, our *in vivo* experiments with EcN demonstrate that bARG_Ser_ expression enables stable, nondestructive acoustic visualization of the microscale distribution of these probiotic agents in a therapeutically relevant context.

### The A. flos-aquae GV gene cluster produces robust nonlinear ultrasound contrast in mammalian cells

Having developed second-generation ARGs for use in bacteria, we attempted the same for mammalian cells. The first-generation mammalian ARGs were based on the GV gene cluster from *B. megaterium* (referred to here as mARG_Mega_). mARG_Mega_ expression could only be detected with destructive collapse-based imaging due to a low level of GV expression and the lack of nonlinear contrast from the resulting GVs.^7^ Moreover, successful use of mARG_Mega_ as a reporter gene required monoclonal selection of transduced cells and their treatment with a broadly-acting histone deacetylase inhibitor^7^. Seeking mARGs that are expressed more robustly and produce nonlinear signal, we cloned mammalian versions of the genes contained in each of the three clusters that produced nonlinear signal in *E. coli* at 37°C: *Serratia, A. flos-aquae,* and *A. flos-aquae/B. megaterium* (**Fig. 1d-e**). Equimolar transient cotransfections of the monocistronic genes derived from each gene cluster into HEK293T cells yielded detectable BURST signal only for the *A. flos-aquae* gene cluster and the positive control mARG_Mega_^7^ (**Fig. 5a, 5b** “1-fold excess”).

**Figure 5.**
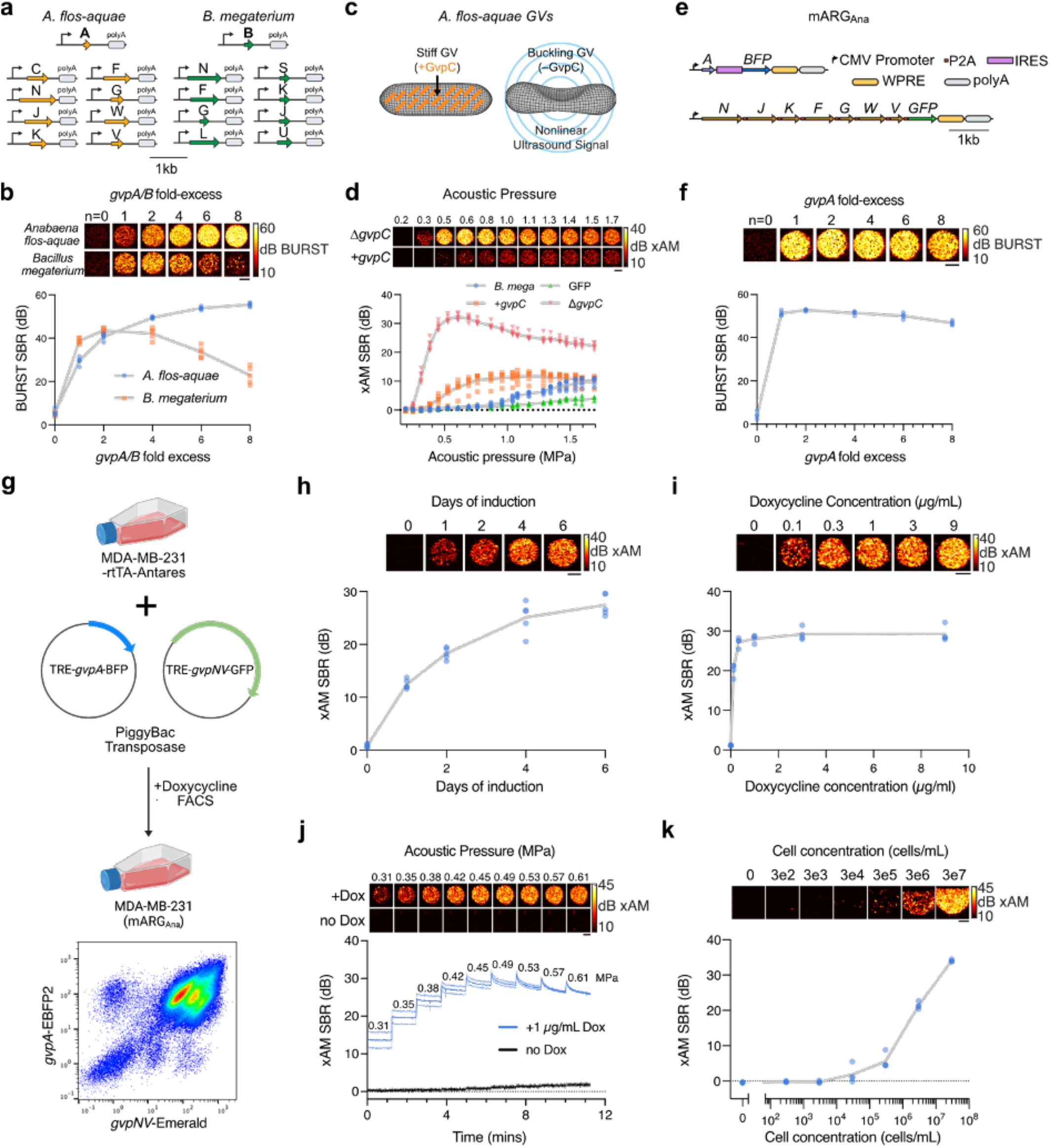
Heterologous expression of the *A. flos-aquae* GV gene cluster in mammalian cells. (**a**) Schematic of the codon-optimized monocistronic plasmid sets used in this study. (**b**) Representative BURST images (top) and SBR quantification (n=5, bottom) of transient GV expression in HEK293T cells 3 days after cotransfection of mixtures with varying *gvpA/B* fold excess relative to their respective assembly factor plasmids. (**c**) Diagram of GV structure with GvpC highlighted in orange. (**d**) Representative xAM ultrasound images (top) and SBR quantification (n=6, bottom) of transient cotransfection experiments *of A. flos-aquae* GV plasmids (4-fold *gvpA* excess) with and without *gvpC* at varying acoustic pressures. *B. megaterium* GV (at 2-fold *gvpB* excess) and GFP expression is included for quantitative comparison. (**e**) Schematic of the mARG_Ana_ plasmid set with fluorescent reporters. (**f**) Representative BURST images (top) and SBR quantification (n=4, bottom) of transient GV expression in HEK293T cells 3 days after cotransfection of mARG_Ana_ mixtures with varying *gvpA* fold excess relative to the assembly factor plasmid. (**g**) Schematic of MDA-MB-231-mARG_Ana_ engineering (created with BioRender.com and FlowJo). The final population was ~95% double positive for *gvpA* and *gvpNJKFGWV* expression. (**h**) Representative xAM images (top) and SBR quantification (n=5, bottom) of MDA-MB-231-mARG_Ana_ cells at 0.54 MPa after 1, 2, 4 and 6 days of 1 μg/ml doxycycline induction. (**i**) Representative xAM images (top) and SBR quantification (n=4, bottom) of MDA-MB-231-mARG_Ana_ cells at 0.42 MPa as a function of doxycycline concentration after 4 days of expression. (**j**) Representative xAM images (top) and SBR quantification (n=4, bottom) of induced and uninduced MDA-MB-231-mARG_Ana_ cells as a function of time under varying acoustic pressures. For j, thick lines represent the mean of 4 replicates and thin lines represent ± standard deviation. For b, d, f, h, and i, gray lines connect the means of the replicates. All ultrasound image scalebars represent 1 mm.

Given the multiple *gvpA* copies contained in the native *A. flos-aquae* GV operon,^8^ we hypothesized that expressing the major structural protein GvpA at a higher stoichiometry relative to the other genes in this cluster could improve GV expression. To test this possibility, we titrated the amount of *gvpA* plasmid in the *A. flos-aquae* plasmid set, while keeping the DNA amount corresponding to other genes constant (the total DNA level was also kept constant with a padding vector). We found that the BURST signal increased monotonically with increasing *gvpA* up to 8-fold *gvpA* excess (**Fig. 5b**). In contrast, the signal peaked at 2-fold excess of the homologous structural protein *gvpB* when expressing the *B. megaterium* cluster, possibly due to unincorporated or unchaperoned GvpB monomers, which may burden the cells. That this does not happen with the genes from *A. flos-*aquae within the range tested suggests that the assembly factors in this gene cluster may be more efficient at utilizing GvpA to form gas vesicles or that GvpA may pose less of a burden than GvpB. To further improve GvpA expression, we stabilized the *gvpA* transcript with WPRE-hGH poly(A) elements,^57^ which resulted in peak signal at lower *gvpA*:chaperone ratios (**Fig. S18, a-b**).

We next looked for nonlinear ultrasound contrast from *A. flos-aquae* GVs to enable their discrimination from background. GvpC is a minor structural protein in *A. flos-aquae* GVs that binds to and mechanically reinforces the GV shell (**Fig. 5c**).^8^ Previously, *in vitro* chemical removal of GvpC from purified GVs was shown to enhance nonlinear ultrasound scattering by allowing the GVs to deform more strongly in response to acoustic pressure.^29,58,59^ When we omitted *gvpC* from our mammalian co-transfection mixture, we observed a dramatic enhancement of nonlinear signal in xAM imaging, with the peak signal achieved at around 0.6 MPa (**Fig. 5d**). By comparison, transfections including *gvpC* produced a much weaker xAM signal, while *B. megaterium* plasmids and GFP-expressing cells did not produce appreciable nonlinear contrast at any pressure. The omission of *gvpC* did not appreciably alter BURST contrast (**Fig. S18c**). These results indicate that the mammalian GVs derived from *A. flos-aquae* can provide strong, nondestructive, nonlinear ultrasound contrast.

To create a convenient vector for mammalian expression of *A. flos-aquae* GVs, we constructed a polycistronic plasmid linking the assembly factor genes *gvpNJKFGWV* through P2A co-translational cleavage elements. *gvpA* was supplied on a separate plasmid to enable stoichiometric tuning. The *gvpA* and *gvpNJKFGWV* plasmids were labeled with IRES-BFP and P2A-GFP, respectively, to allow for fluorescent analysis and sorting. Both transcripts were driven by CMV promoters and stabilized by WPRE-hGH poly(A) elements. We termed this pair of plasmids mARG_Ana_: mammalian ARGs adapted from *A. flos-aquae* (**Fig. 5e**). mARG_Ana_ produced robust GV expression and ultrasound contrast in HEK293T cells transiently co-transfected with a 1-to 6-fold molar excess of *gvpA* (**Fig. 5f**).

One of the most promising applications of mammalian reporter genes is the visualization of tumor growth in animal models of cancer, which are a critical platform for basic oncology research and the development of new therapeutics. To produce a stable cancer cell line expressing mARG_Ana_, we cloned our polycistronic constructs into PiggyBac integration plasmids under a doxycycline-inducible TRE promoter.^60,61^ As a clinically relevant cancer model, we chose the human breast cancer cell line MDA-MB-231, which is widely used in tumor xenograft studies. We engineered these cells to constitutively co-express the rtTA transactivator and Antares optical reporter^62^, then electrically transduced them with a mixture of mARG_Ana_ and PiggyBac transposase plasmids at a 2:1 *(gvpA:gvpNJKFGWV)* molar ratio, and fluorescently sorted for co-expression of Antares, GFP and BFP (**Fig 5g** and **Fig. S18d**).

The resulting polyclonal MDA-MB-231-mARG_Ana_ cells showed xAM contrast after a single day of doxycycline induction, which increased substantially through day 6 (**Fig. 5h**). We confirmed the expression of GVs in these cells by electron microscopy (**Fig. S18e**). The ultrasound signal increased steeply with increasing doxycycline doses up to 1 μg/mL, above which the signal saturated (**Fig. 5i**). xAM signal was detected from induced cells starting from an acoustic pressure of 0.31 MPa, whereas uninduced control cells did not produce signal at any pressure (**Fig. 5j**). At pressures above 0.42 kPa, the xAM signal gradually decreased over time, indicating the partial collapse of GVs. We chose 0.42 MPa as the xAM imaging pressure for subsequent experiments, providing the optimal balance of signal stability and signal strength.

To test whether second-generation mammalian ARGs generalize beyond HEK293T and MDA-MB-231 cells, we also engineered mouse 3T3 fibroblasts and human HuH7 hepatocytes to express the mARG_Ana_ operon. Both cell types showed excellent xAM contrast, providing a similar xAM detection sensitivity as MDA cells (**Fig. 5k, Fig. S18f**). At 300k cells/mL for MDA-MB-231 and 3T3, and at 30k cells/mL for HuH7, the detection limit surpassed the reported destructive imaging sensitivity of first-generation mARG_Mega_ by one and two orders of magnitude respectively.^7^ With BURST imaging, mARG_Ana_ cells could be detected still more sensitively, at concentrations down to 3,000 cells/mL for 3T3-mARG_Ana_ and 30,000 cells/mL for MDA-MB-231-mARG_Ana_ and HuH7-mARG Ana **(Fig. S18g)**.

### mARG_Ana_ expression enables visualization of in vivo gene expression patterns in an orthotopic tumor model

We next tested the ability of mARG_Ana_ to reveal the spatial distribution of gene expression in tumor xenografts in living mice. We formed orthotopic tumors by injecting MDA-MB-231-mARG_Ana_ cells bilaterally in the fourth mammary fat pads of female immunocompromised mice. The mice were then split into doxycycline-induced and uninduced groups. We acquired ultrasound images of the tumors as they grew, with 3 imaging sessions distributed over 12 days (**Fig. 6a**). All induced tumors produced bright and specific xAM contrast starting from the first timepoint (day 4), whereas the uninduced tumors did not (**Fig. 6b**). The acquisition of adjacent planes allowed 3D visualization of expression patterns (**Supplementary Video 1**). The nonlinear xAM signal was highly specific to the viable tumor cells, being absent outside the anatomically visible tumor boundaries and within the cores of the larger tumors imaged at 8 and 12 days. The observed spatial pattern of gene expression in these tumors was corroborated by fluorescence microscopy of formalin-fixed tumor sections obtained from euthanized mice on day 12 (**Fig. 6c**, **Fig. S19**), confirming the ability of mARG_Ana_ to report microscale patterns of gene expression noninvasively in living animals. In contrast, *in vivo* fluorescence images of the mice lacked information about the spatial distribution of gene expression within the tumor (**Fig. 6d**). We quantified GV expression over time as a volumetric sum of xAM signal over all acquired image planes (**Fig. 6e**). The induced tumors had significantly higher total signal than the uninduced controls at all time points. These experiments demonstrate the ability of mARG_Ana_ to serve as a highly effective reporter gene for the noninvasive monitoring of tumor growth and gene expression.

**Figure 6.**
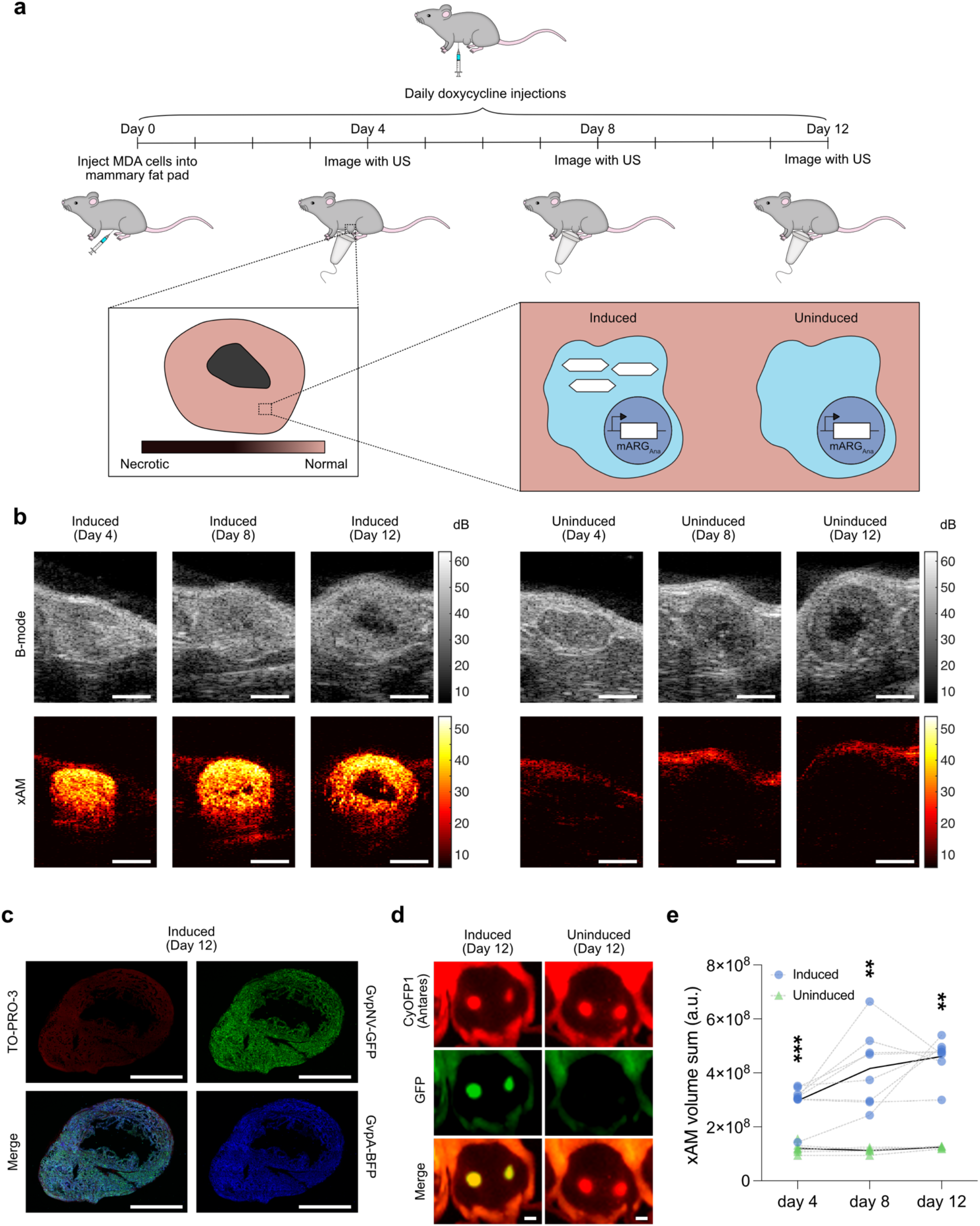
*In situ* mARG_Ana_ expression enables nondestructive ultrasound imaging of orthotopic tumors. (**a**) Diagram of the *in vivo* protocol for assessing *in situ* mARG_Ana_ expression in orthotopic tumors. Mice were injected bilaterally in 4^th^ mammary fat pads with engineered MDA-MB-231-mARG_Ana_ human breast adenocarcinoma cells on day 0. mARG_Ana_ expression was induced by regular intraperitoneal (IP) doxycycline injections starting from the day of tumor injections. Tumors were imaged with ultrasound after 4, 8 and 12 days of expression. (**b**) Representative middle sections of B-mode and xAM ultrasound tomograms of MDA-MB-231 mARG_Ana_ tumors induced with doxycycline (left) and uninduced control (right) imaged on day 4, 8 and 12. Scalebars represent 2 mm. See supplementary video for the full ultrasound tomogram of the induced tumor at 12 days. (**c**) Fluorescence micrograph of a 100 nm thin tumor section. Green color shows GFP fluorescence, blue color shows BFP fluorescence, and red color shows TO-PRO-3 nuclear stain. See **Fig. S17** for the uninduced control. Scalebars represent 2 mm. (**d**) Whole-animal fluorescence imaging of induced (left) and uninduced (right) mouse after 12-days of expression. All tumors are constitutively expressing CyOFP1 (Antares, red) whereas mARG_Ana_ expression is linked to expression of GFP (green). The left (reader’s right) tumors are shown in panel (b). Scalebars represent 5 mm. (**e**) Three-dimensional sum of xAM signal from ultrasound tomograms of induced (n=8) and uninduced (n=7 on day 4, n=5 on days 8 and 12) tumors from all three imaging sessions plotted on a linear scale in arbitrary units. Asterisks represent statistical significance by unpaired t tests between induced and uninduced conditions (*** = p<0.001, ** = p<0.01, * = p<0.05, ns = no significance).

The primary advantage of ultrasound imaging over optical methods is deeper penetration. To ensure that mARG_Ana_ expression can be detected in deep tissue beyond what can be easily shown in mice, we imaged MDA-MB-231-mARG_Ana_ cells using xAM through a slab of beef liver thicker than 1 cm. As expected, the MDA-MB-231-mARG_Ana_ cells were readily detectable (**Fig. S19b**). Moreover, this xAM signal was highly specific to MDA-MB-231-mARG_Ana_ cells, and did not appear in the ARG-negative liver tissue (**Fig. S19c**).

### Real-time nondestructive ultrasound imaging of mARG_Ana_ expression enables ultrasound-guided biopsy of genetically defined cell populations

One of the most common uses of ultrasound in biological research and medicine is to spatially guide procedures such as biopsies. By enabling non-destructive imaging, second-generation mARGs create the possibility for such procedures to be targeted based on *in vivo* gene expression patterns. Notably, because real-time procedure guidance requires dynamic non-destructive imaging, such guidance was not possible with previous ARGs. To demonstrate this concept, we performed ultrasound-guided biopsies on chimeric tumors created by making adjacent subcutaneous injections of MDA-MB-231-mARG_Ana_ cells and MDA-MB-231 cells expressing Antares (**Fig. 7a**). We visualized the chimeric composition of these tumors by obtaining 3D tomograms of the tumor mass (**Supplementary Video 2, 3).** We then performed fine needle aspiration biopsies targeting either the xAM-positive or the xAM-negative regions of each tumor (**Fig. 7b, Supplementary video 4, 5**). Flow cytometric analysis of dissociated biopsy samples showed a very high correlation between mARG_Ana_-positive (GFP-positive) cells that were sampled from xAM-positive regions and vice versa (**Fig. 7c, Fig. S20)**. These results demonstrate that mARG_Ana_ expression enables genetically targeted *in vivo* procedures under ultrasound guidance.

**Figure 7.**
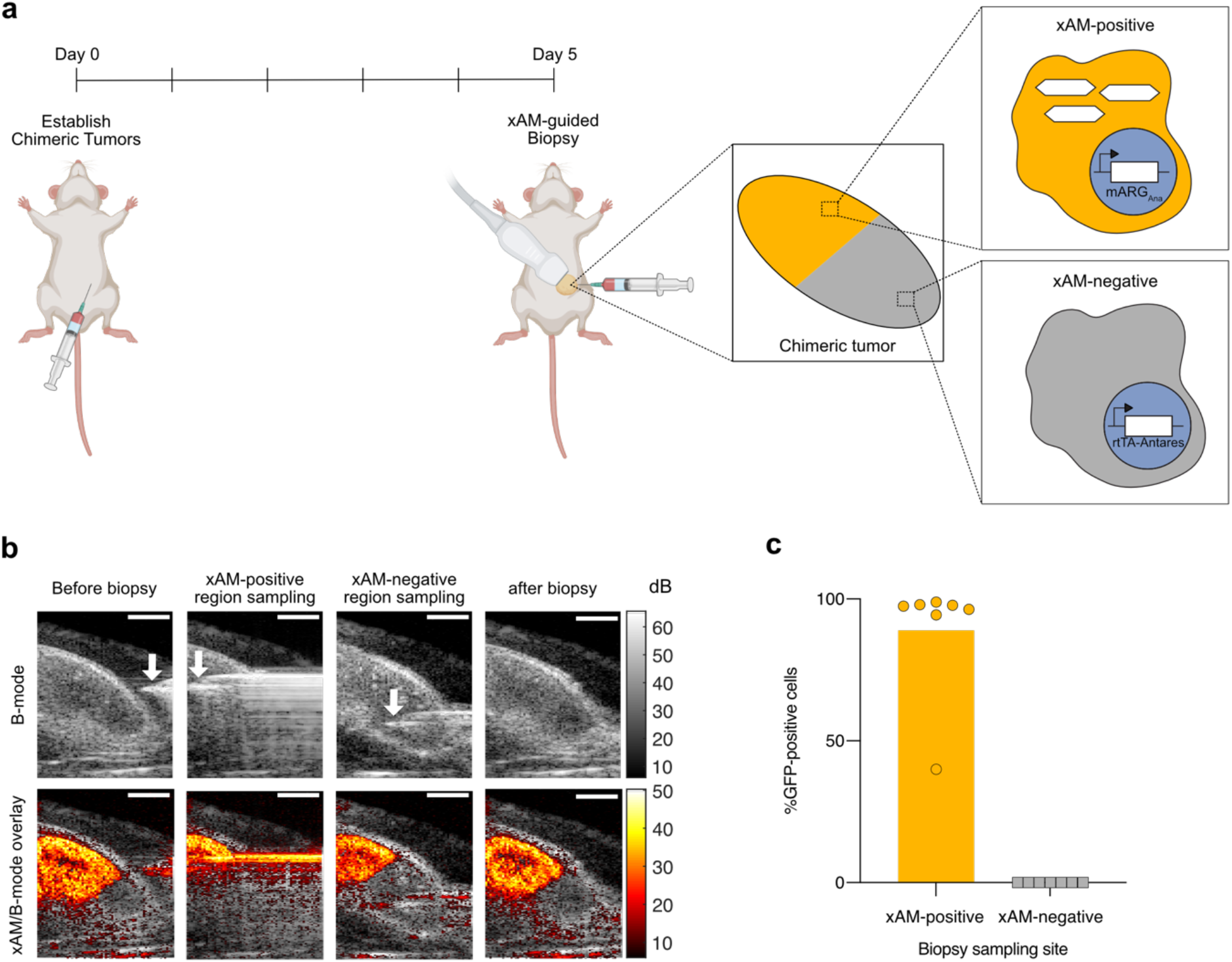
xAM imaging of mARG_Ana_ enables ultrasound-guided genetically selective tumor biopsy. (**a**) Diagram of the *in vivo* protocol for establishing chimeric tumors, *in situ* expression of GVs and tumor biopsy. Chimeric tumors were established on day 0. mARG_Ana_ expression was induced by regular intraperitoneal (IP) doxycycline injections starting from the day of tumor injections. Tumors were imaged with ultrasound and biopsied after 5 days of expression. (**b**) Representative B-mode and xAM an ultrasound-guided biopsy procedure. Scalebars represent 2 mm. See Supplementary Videos 2,3,4 and 5 for the full ultrasound tomogram of the induced chimeric tumor, 3D reconstruction of the chimeric tumor as well as the videos of the full procedure. (**c**) Results of flow cytometric analysis of biopsied samples from xAM-positive and xAM-negative regions of chimeric tumors (n=7 tumors). See **Fig. S20** for flow cytometric gating strategy. Bar height represents the mean, circles represent individual data points.

## DISCUSSION

Our results establish two second-generation ARG constructs bARG_Ser_ and mARG_Ana_ that provide unprecedented, realtime ultrasound detection sensitivity and specificity when expressed in bacteria and mammalian cells. These gene clusters, obtained through a systematic phylogenetic screen and optimized through genetic engineering, produce bright nonlinear ultrasound signal when expressed *in situ* in tumors, either by bacterial agents colonizing the necrotic core of a tumor, or by the tumor cells themselves. When imaged using a highly sensitive and specific non-destructive ultrasound imaging paradigm, this nonlinear signal enables real-time monitoring of the precise locations and transcriptional activities of these cells and is sufficiently stable to image cellular biodistribution and gene expression over multiple days. Furthermore, real-time nondestructive imaging of ARGs enables ultrasound-guided procedures such as biopsies to target genetically defined cells.

These results comprise a major advance over previous work on heterologous GV expression in bacterial and mammalian cells. Previous bARG constructs required the host bacteria to be cultured and pre-express GVs under ideal laboratory conditions before *in vivo* injection,^6^ while the previous mARG construct only produced ultrasound contrast when expressed in a sorted monoclonal cell line treated with a global epigenetic activator.^7^ In both cases, sensitive and specific detection of these cells relied on destructive imaging methods that produce one-time contrast, hindering their use in monitoring dynamic biological processes or guiding real-time interventions.^10^

With these major improvements, we anticipate that these new ARGs will be useful for many applications that demand the noninvasive imaging of cells deep inside the body. bARGs could be used to track therapeutic bacteria as they home to and proliferate in tumors or other target organs. The distribution of the bacteria in the tumor provides a critical readout of whether the therapy is working as designed or needs to be adjusted or re-administered (for example, when the tumor is only partially colonized).^63–66^ In addition, the spatial distribution is important in guiding focused energy procedures applied to the cells, such as focused ultrasound.^67–69^ Other potential applications include the study of the gastrointestinal (GI) microbiome and the tracking of probiotics designed to diagnose or treat GI conditions.^70^,^71^ In these applications, luminescence imaging would provide limited *in vivo* resolution due to light scattering, and would be difficult to scale to larger animals or human patients.

Second-generation mARGs could be used in biological research to visualize the growth and viability of tissues such as tumors and sample their contents with ultrasound-guided procedures. Similar approaches could be applied to study the immune system, the brain and organismal development, where the mARGs could be expressed constitutively or under phenotype-dependent promoters. In addition, mARG expression could enable the tracking of therapeutic mammalian cells, such as T-cells, stem cells and transplants analogously to the imaging of bacterial agents demonstrated in this study, and provide a means to target these therapeutic cells for biopsies and other image-guided interventions. Moreover, the fact that the new mARGs are based on *A. flos-aquae* GVs creates a connection between mammalian expression and molecular engineering 2,6,58,72, including the development of acoustic biosensors to monitor more dynamic cellular signals such as enzyme activity.^72^

The envisioned applications of both bacterial and mammalian ARGs will benefit from the relative simplicity and low cost of ultrasound compared to other non-invasive techniques such as nuclear imaging and MRI, while providing *in vivo* resolution and potential for human translatability beyond what is currently possible with optical methods.

While these results represent an important step in the development of ARGs, additional improvements could further expand their utility in biotechnology. First, the expression kinetics of ARGs are slower than those of fluorescent proteins in bacteria^73^ and mammalian cells,^74^ and faster expression would facilitate the imaging of more dynamic genetic outputs. However, there are numerous scenarios in which expression kinetics on the order of one day are acceptable. For example, the processes of tumor growth and tumor infiltration by therapeutic cells are adequately captured, as shown in this study. Additional examples include inflammation, mammalian development, and stem cell expansion and migration. To enable a broader range of applications, the kinetics of ARG expression could potentially be accelerated through further genetic engineering, for example by pre-expressing certain genes in the ARG cluster *(e.g.,* the assembly factors) and conditionally expressing the remaining genes (*e.g.*, the structural proteins). Second, the ability to multiplex different “colors” of ARGs nondestructively *in vivo* would enable discrimination between different strains of engineered bacteria in a consortium,^75–77^ between different mammalian cell types in a tissue, or between bacterial and mammalian cells in close proximity, such as in a bacterially-colonized tumor. Third, the *Serratia* GV gene cluster is relatively large, making it more challenging to clone and incorporate with other genetic elements. The engineering of a shorter cluster with similar acoustic properties would simplify these efforts. Similarly, the current mARG_Ana_ cluster is delivered using two polycistronic plasmids; consolidating this cluster into a single plasmid would increase transfection efficiency, and shortening it could facilitate its viral packaging and delivery to endogenous cells *in situ*. Fourth, the potentially burdensome effects of large plasmid size and ARG expression on cellular metabolism may be more pronounced *in vivo* than *in vitro* under ideal growth and expression conditions (**Figs. 2, 5h-i, S8–11**). While the *in vivo* expression of both bARG_Ser_ (**Fig. 4)** and mARG_Ana_ (**Fig. 6**) yielded strong and specific ultrasound signals without obvious effects on cell migration and growth, strategies to reduce construct size and tightly regulate ARG expression could become more important as the range of ARG applications expands. Finally, while mice are the predominant animal model in biomedical research and an established proving ground for reporter gene technologies^78–82^, future work is needed to test ARGs in additional species. The ability of second-generation ARGs, demonstrated here, to function in cells from two different mammalian species and two different bacterial species, is an encouraging sign for their broader utility.

Our phylogenetic screening approach was successful in identifying bARG_Ser_ and mARG_Ana_ as greatly improved ARGs. However, out of practical necessity, this screen subsampled the available phylogenetic space, and testing additional GV-encoding gene clusters could reveal ARGs with new or further-improved properties. Improvements can also be made in the screening strategy. It is challenging to identify all the *gvp* genes in a given genome (see Methods), and there is considerable regulation of GV cluster transcription by factors inside^9,32^ or outside^33^ the clusters. Therefore, in a given cloned cluster, it is possible that either essential genes are missing or cryptic regulatory elements are included.^9,30,34^ These issues could be resolved by synthesizing multiple versions of each putative gene cluster and screening a larger number of them in higher throughput. Even with optimal genetic constructs, it is likely that gene clusters from some species will not successfully form GVs in a given heterologous host due to differences in growth temperature, turgor pressure or the presence or absence of specific host factors. The phylogenetic screening strategy used in this study could thus be adapted to find optimal ARGs for use in other species of interest.

Just as improvements to and adaptations of fluorescent proteins enabled a wide range of microscopy applications that were mere speculations when GFP was first harnessed as a biotechnology, the systematic development of next-generation ARGs will help bring to reality the promise of sensitive, high-resolution noninvasive imaging of cellular function inside intact mammals.

## MATERIALS AND METHODS

### Genomic mining of ARG clusters

A literature search was conducted to find papers reporting the production of gas vesicles in any species. Search terms included “gas vesicle,” “gas vacuole,” “aerotope,” and “aerotype.” All species found are listed in **Table S1**. If the report did not include a strain name, then any available 16S rRNA gene sequence was used (as it was assumed that any other strain of the same species would fall in the same place on the phylogenetic tree), but no GV gene cluster sequence was used (even if it was available for one or more strains of that species) because it was found during our analysis that: 1) several reports describe species for which some strains produce GVs but others don’t, and 2) comparison of GV gene cluster sequences of multiple strains of the same species almost always showed differences — often very significant ones. Further, even if a reference stating that a given organism produced GVs was not available, 16S rRNA gene sequences from all members of the following genera were included because GV production is a diacritical taxonomic feature for these genera: *Dolichospermum*,^84^ *Limnoraphis^85^* and *Planktothrix*.^86^

GV clusters were identified in genomes through a combination of annotations and sequence similarity to known *gvp* genes. However, there were two challenges in identifying all *gvp*s in a given genome: 1) there is little to no annotation for many *gvps,* and 2) GV gene clusters are not always contiguous in genomes, and *gvp*s can occasionally be found hundreds of kb away from the main cluster(s). We attempted to only select “well-behaved” GV clusters for testing *(i.e.,* ones in which all *gvp*s identified in that genome were present in clusters, and these clusters contained a minimum of non-*gvp* genes, which could increase the metabolic burden of cluster expression without increasing GV yield), but it is possible that even for these clusters, some *gvp*s were not cloned.

Of our list of 288 strains reported to form gas vesicles, 270 had 16S rRNA gene sequences available (**Table S1**). These were downloaded from NCBI using a custom Python script, and a multiple sequence alignment was constructed using Clustal Omega.^87^ This alignment was used to generate a phylogenetic tree file using ClustalW2,^88^ which was rendered using EvolView.^89^ Only unique species are displayed in the phylogenetic trees in **Fig. 1a** and **Fig. S1**.

### Bacterial plasmid construction and molecular biology

Organisms were obtained from commercial distributors as indicated in **Table S2**. If an organism was shipped as a liquid culture, the culture was centrifuged and the pellet resuspended in ddH2O, as it was found that even trace amounts of certain culture media could completely inhibit PCR. Fragments were amplified by PCR using Q5 polymerase and assembled into a pET28a(+) vector (Novagen) via Gibson Assembly using reagents from New England Biolabs (NEB). Sub-cloning and other modifications to plasmids were also performed with Gibson Assembly using reagents from NEB. Assemblies were transformed into NEB Stable *E. coli*. All constructs were verified by Sanger sequencing.

*Halobacterium salinarum* has two chromosomal GV gene clusters (plus additional plasmid-borne ones), which were cloned and tested separately. *Methanosarcina vacuolata* has only one cluster, but while its genome sequence in the NCBI database has two copies of *GvpA1* and one copy of *GvpA2,* our genomic PCR yielded a product with only one copy of *GvpA1.* In a second cloning step, we added a copy of *GvpA2* to the cloned cluster. While we were able to PCR *GvpA2* from the genome, it was not contiguous with the rest of the cluster. Therefore, we speculate that either there was an error in the assembly of the genome sequence (likely caused by the high sequence similarity of the *GvpA* genes), or that the genotype of our strain differs slightly from that of the strain sequenced.

### In vitro bacterial expression of ARGs

For initial testing, all constructs were expressed in BL21(DE3) *E. coli* (NEB). Fifty μL of electrocompetent *E. coli* were transformed with 1.5 μL of purified plasmid DNA (Econospin 96-well filter plate, Epoch Life Science), and 1 mL of SOC medium (NEB) was added immediately after electroporation. These cultures were incubated at 37°C for 2 hr, and 150 uL was inoculated into larger 1.5 mL LB cultures containing 100 ug/mL kanamycin and 1% (w/v) glucose (for catabolite repression^90^ of the BL21(DE3) PlacUV5 promoter) in a deepwell 96-well plate and grown overnight in a shaking incubator at 30°C. Square dual-layer LB agar plates were prepared as described previously,^6^ with varying concentrations of IPTG and 100 ug/mL kanamycin in the bottom layer, and 1% (w/v) glucose and 100 ug/mL kanamycin in the top layer. LB agar was incubated at 60°C for 12-36 hr after dissolution to allow it to degas. After the agar solidified, plates were dried at 37°C to remove all condensation on the top layer that would cause the bacterial patches to run together. A multichannel pipette was used to thoroughly mix overnight cultures and drop 1 μL of each culture onto the surface of the dual-layer plates, with care taken to avoid puncturing the agar which results in artifacts during ultrasound scans. Importantly, low-retention pipette tips were used, as it was found that the small volumes of culture would wet the outsides of standard pipette tips, resulting in inaccurate volume dispensing. Patches were allowed to dry completely before plates were flipped and incubated at 37°C for 24 hr or 30°C for 48 hr.

For *in vitro* expression experiments in EcN, the appropriate plasmids were first transformed via electroporation and the outgrowth was plated on LB (Miller)-agar plates with the appropriate antibiotic (25 μg/mL chloramphenicol or 50 μg/mL kanamycin) and 1% (w/v) glucose. The resulting colonies were used to inoculate 2 mL LB (Miller) medium with the appropriate antibiotic and 1% (w/v) glucose, and these cultures were incubated at 250 rpm and 37°C overnight. Glycerol stocks were prepared by mixing the overnight cultures in a 1:1 volume ratio with 50% (v/v) glycerol and storing at −80°C. The night before expression experiments, glycerol stocks were used to inoculate overnight cultures (2 mL LB medium with the appropriate antibiotic and 1% (w/v) glucose) which were incubated at 37°C and shaken at 250 rpm. For expression on solid media, 1 μL of overnight culture was dropped onto square dual-layer LB agar plates with 2X the final inducer (IPTG, aTc, or L-arabinose) concentration in the bottom layer, 1% (w/v) glucose in the top layer, and the appropriate antibiotic in both layers (50 μg/mL chloramphenicol or 100 μg/mL kanamycin). Plates were allowed to dry, and then inverted and incubated at 37°C for 24 hours before imaging with ultrasound. For expression in liquid media, 500 μL of each overnight culture was used to inoculate 50 mL LB supplemented with 0.4% (w/v) glucose and 25 μg/mL chloramphenicol in 250 mL baffled flasks. Cultures were incubated at 37°C and 250 rpm until reaching at OD_600_ of 0.1 - 0.3. At this point, cultures were induced by addition of L-arabinose and placed back at 37°C and 250 rpm. For time titration experiments, 0.1% (w/v) L-arabinose was used for induction and 0.5 mL of each culture was removed at 0, 1, 3, 5, 7, 9, 11, and 24 hours post-induction for OD_600_ and ultrasound measurements. For L-arabinose titration experiments, L-arabinose concentrations ranging from 0 to 1% (w/v) were used for induction, and cultures were incubated for 24 hours at 37°C and 250 rpm after addition of L-arabinose before ultrasound imaging. For cell concentration titration experiments, cultures were incubated for 24 hours at 37°C and 250 rpm after addition of 0.1% (w/v) L-arabinose before ultrasound imaging. All cultures were stored at 4°C or on ice until casting in phantoms and imaging with ultrasound. In all liquid culture experiments, cultures were prescreened for the presence of GVs by phase contrast microscopy before being imaged with ultrasound.

*In vitro* expression experiments in *S. Typhimurium* were performed as described above for EcN, except plasmids were transformed into an attenuated version of *Salmonella enterica* serovar Typhimurium strain SL1344^50^, 2xYT medium was used instead of LB medium, and induction in liquid culture was performed by adding L-arabinose to the medium at the time of inoculation at an OD_600_ of 0.05.

To assess plasmid stability of pBAD-bARG_Ser_-AxeTxe in EcN, the glycerol stock of this strain was used to inoculate 2 mL LB (Miller) supplemented with 25 μg/mL chloramphenicol and 1% (w/v) glucose, and this culture was incubated at 37°C and 250 rpm overnight. Twenty μL of the overnight culture was subcultured into 2 mL LB with 25 μg/mL chloramphenicol, 2 mL LB without antibiotics, and 2 mL LB without antibiotics and with 0.1% (w/v) L-arabinose, each in quadruplicate. Every 24 hours, 20 μL of each culture was sub-cultured into fresh media of the same conditions. All cultures were incubated at 37°C and 250 rpm. On days 1-3, 5, and 7, serial dilutions of each culture were plated on LB-agar without antibiotics, LB-agar with 25 μg/mL chloramphenicol, and LB-agar with 25 μg/mL chloramphenicol + 0.1% (w/v) L-arabinose + 0.4% (w/v) glucose. Plates were incubated at 37°C for at least 16 hours and colonies were counted and screened manually. Plasmid retention was assessed by taking the ratio of CFUs on LB-agar plates with chloramphenicol to CFUs on LB-agar plates without antibiotics. The presence of mutations that disrupt the ability to express functional bARG_Ser_ was assessed by a loss of colony opacity on LB-agar plates with 25 μg/mL chloramphenicol + 0.1% (w/v) L-arabinose + 0.4% (w/v) glucose.

### In vitro ultrasound imaging of bacteria expressing ARGs on solid media

Ultrasound imaging of bacterial patches was performed using a Verasonics Vantage programmable ultrasound scanning system and an L10-4v 128-element linear array transducer (Verasonics) with a center frequency of 6 MHz and an element pitch of 300 μm. Image acquisition was performed using a custom imaging script with a 64-ray-lines protocol and a synthetic aperture of 65 elements. The transmit waveform was set to a voltage of 50 V and a frequency of 10 MHz, with 1 waveform cycle and 67% intra-pulse duty cycle. In xAM mode, a custom sequence detailed previously^29^ was used with an angle of 19.5°. RF data from 4 repeated acquisitions was coherently averaged prior to beamforming for each image plane.

Agar plates containing bacterial patches were coated with a thin layer of LB agar and immersed in PBS to allow acoustic coupling to the L10-4v transducer. The transducer was connected to a BiSlide computer-controlled 3D translatable stage (Velmex) and positioned above the plane of the plate at an angle of 15° from the vertical (to minimize specular reflection from the plastic dishes and agar) and a distance of 20 mm from the bacterial patches. The imaging sequence was applied sequentially to acquire image planes covering the full area of all plates. A custom script was used to automate the scan by controlling the motor stage in tandem with the ultrasound system, translating 0.5 mm in the azimuthal direction between rows and 19.5 mm in the lateral direction between columns. In the case of differential imaging scans, the full scan sequence was repeated after returning the motor stage to its origin and adjusting the voltage of the transducer.

For image processing and analysis, custom beamforming scripts were applied on-line to reconstruct image planes from the acquired RF data at each location. The intensity data for each plane was saved for off-line processing. All image planes were concatenated to form a 3D volume with all plates and colonies. A 2D image of the colonies was extracted from the 3D volume by taking the maximum intensity over a manually-defined depth range for all voxel columns. 2D differential images were obtained by subtracting the post-collapse 2D image from the pre-collapse 2D image. Bacterial patch intensities were then quantified from these 2D images. Sample ROIs were drawn around the center of each patch to avoid artefacts from the edges, and background ROIs were drawn around representative regions without patches. The signal-to-background ratio (SBR) was calculated as the mean pixel intensity of the sample ROI divided by the mean pixel intensity of the background. Conversion to decibels (dB) was calculated as 20*log10(SBR). For display, images were normalized by dividing by the average background signal of all images being compared and setting the lower and upper limits of the colormaps to be the same, where the lower limit was equal to a constant A times the average background and the upper limit was equal to a constant B times the maximum pixel intensity out of all images being compared; images were then converted to dB. For xAM and differential xAM images of bacterial patches, A was set to 1 and B was set to 0.5.

### In vitro ultrasound imaging of bacteria expressing ARGs suspended in agarose phantoms

To create phantoms for ultrasound imaging of bacteria from liquid cultures or suspended in PBS from patches on solid media, wells were cast with a custom 3D-printed mold using 1% (w/v) agarose in PBS, which was degassed by incubating at 65°C for at least 16 hours. Cultures or cell suspensions to be analyzed were diluted in ice-cold PBS to 2x the final desired cell concentration (calculated from the measured OD_600_), incubated at 42°C for one minute, and mixed 1: 1 with 1 % (w/v) agarose in PBS at 42°C for a final concentration of 1x. This mixture was then loaded into the wells in duplicate and allowed to solidify. Care was taken not to introduce bubbles during this process. The phantoms were submerged in PBS, and ultrasound images were acquired using a Verasonics Vantage programmable ultrasound scanning system and an L22-14v 128-element linear array transducer with a center frequency of 18.5 MHz with 67%-6-dB bandwidth, an element pitch of 100 μm, an elevation focus of 8 mm, and an elevation aperture of 1.5 mm. The transducer was attached to a custom-made manual translation stage to move between samples. B-mode and xAM images were acquired using the same parameters as described previously:^72^ the frequency and transmit focus were set to 15.625 MHz and 5 mm, respectively, and each image was an average of 50 accumulations. B-mode imaging was performed with a conventional 128-ray-lines protocol, where each ray line was a single pulse transmitted with an aperture of 40 elements. xAM imaging was performed using a custom sequence detailed previously^29^ with an angle of 19.5° and an aperture of 65 elements. The transmitted pressure at the focus was calibrated using a Fibre-Optic Hydrophone (Precision Acoustics), and the peak positive pressure was used as the “acoustic pressure” in Fig. 3. BURST images were acquired as a series of pAM images as described previously,^7^ except the focus was set to 6 mm, and the acoustic pressure was set to 0.15 MPa (1.6V) for the first 10 frames and 3.0 MPa (25V) for the last 46 frames.

To measure the xAM signal at varying acoustic pressures, an automated voltage ramp imaging script was used to acquire an xAM image at each voltage step (0.5 V increments from 2 to 25 V), immediately followed by a B-mode acquisition at a constant voltage of 1.6 V (0.15 MPa) before another xAM acquisition at the next voltage step; the voltage was held constant for 10 seconds at each step before the image was saved. To measure the xAM and B-mode signals over time at various acoustic pressures, another script was used to automatically save an xAM or B-mode image every second while the voltage was automatically increased by 2 V approximately every 70 seconds. Each frame consisted of 64 ray lines, which took 180 μs each to acquire, giving a pulse repetition rate of 86.8 Hz. Based on these results, all subsequent *in vitro* xAM images of bARG_Ser_-expressing EcN were acquired at 18V (1.29 MPa).

For experiments in Fig. S12 and S14, a different transducer, an L22-14vX transducer, was used which had a different pressure-to-voltage calibration. Consequently, for ultrasound imaging of *S. Typhimurium,* xAM imaging was performed at 1.72 MPa (14V), unless otherwise noted, and BURST was performed using 0.16 MPa (1.6V) for the first 10 frames and 3.7 MPa (25V) for the final 46 frames.

xAM and B-mode image processing and analysis were performed as described above, except that custom beamforming scripts were applied off-line to reconstruct images from the saved RF data for each sample, no 3D reconstruction was performed as images captured at single locations, circular ROIs were drawn around sample and background regions (taking care to avoid bubbles) to calculate SBRs, and values of A= 1.4 and B=0.5 were used to normalize images for display. BURST images were reconstructed using the signal template unmixing algorithm as described previously^35^; as above, circular ROIs were then drawn around sample and background regions to calculate SBRs and values of A=3 and B=1 were used to normalize images for display.

### Microscopy of bacteria

For TEM imaging, cells expressing GVs were diluted to OD_600_ ~1 in 10 mM HEPES (pH 7.5) or culture media. 3 μL of the sample was applied to a freshly glow-discharged (Pelco EasiGlow, 15 mA, 1 min) Formvar/carbon-coated, 200 mesh copper grid (Ted Pella) for 1 min before being reduced to a thin film by blotting. Grids with cells were washed three times in 10 mM HEPES (pH 7.5), blotted, air-dried, and imaged without the stain. Image acquisition was performed using a Tecnai T12 (FEI, now Thermo Fisher Scientific) electron microscope operated at 120 kV, equipped with a Gatan Ultrascan 2k X 2k CCD.

For phase contrast microcopy (PCM) imaging, cells expressing GVs were scraped off from plates and re-suspended in PBS at an OD_600_ of 1-2, or liquid cultures were used directly. Suspensions were transferred to glass slides and PCM images were acquired using a Zeiss Axiocam microscope with a 40X Ph2 objective.

### In vivo bacterial ARG expression and ultrasound imaging

All *in vivo* experiments were performed under a protocol approved by the Institutional Animal Care and Use of Committee (IACUC) of the California Institute of Technology. For experiments involving tumor colonization with EcN, MC26 cells were grown in DMEM media in T225 flasks. After trypsinization and resuspension in PBS + 0.1 mg/mL DNAseI, 5 × 10^6^ MC26 cells were injected subcutaneously into the right flank of 6–8-week-old female Balb/cJ mice. Tumors were allowed to grow for 14 days (reaching sizes of 200-300 mm^3^) before injecting 10^8^ EcN cells suspended in PBS via the lateral tail vein. The day before injection of EcN, Ibuprofen was added to the drinking water at 0.2 mg/mL to ameliorate side effects of EcN injections. To prepare the EcN for injection, the appropriate glycerol stocks were used to inoculate 2 mL LB + 1% (w/v) glucose + 25 ug/mL chloramphenicol which was incubated at 37°C and 250 rpm overnight. The overnight culture (500 μL) was used to inoculate 50 mL LB + 0.4% (w/v) glucose + 25 μg/mL chloramphenicol in 250 mL baffled flasks, which was grown at 37°C and 250 rpm until reaching an OD_600_ of 0.3 - 0.6. This culture was pelleted, washed 4 times with PBS, resuspended in PBS at an OD_600_ of 0.625, and used for injection. Three days after injection of EcN, mice were injected intraperitoneally with 120 mg L-arabinose to induce the EcN. Starting 24 hours after induction, ultrasound images of tumors were acquired as described below. After imaging, 3.0 MPa acoustic pressure was applied throughout the tumor to collapse GVs, and mice were injected again intraperitoneally with 120 mg L-arabinose. The next day, mice were imaged again with ultrasound for re-expression of GVs. The following day, all mice were euthanized and tumors were excised, homogenized, serially diluted, and plated on selective media (LB-agar + 25 μg/mL chloramphenicol) as well as on induction plates (LB-agar + 25 μg/mL chloramphenicol + 0.4% (w/v) glucose + 0.1% (w/v) L-arabinose). Colonies on plates with chloramphenicol were manually counted to quantify the levels of colonization, and colonies on induction plates were screened for a non-opaque mutant phenotype.

For ultrasound imaging, mice were anesthetized with 2% isoflurane and maintained at 37°C using a heating pad. Images were acquired using the L22-14v transducer attached to a manual translation stage described above. Any hair on or around the tumors was removed with Nair, and Aquasonic 100 ultrasound transmission gel was used to couple the transducer to the skin. Parabolic B-mode and parabolic AM (pAM) images were first acquired using a custom 128 ray line script. Each image was formed from 96 focused beam ray lines, each with a 32-element aperture and 6 mm focus. The transmit waveform was set to a voltage of 1.6V in B-mode or 8V in pAM and a frequency of 15.625 MHz, with 1 waveform cycle and 67% intra-pulse duty. In B-mode, each ray line was a single transmit with all 32 elements, and in pAM each ray line consisted of one transmit with all 32 elements followed by 2 transmits in which first the odd and then the even-numbered elements are silenced.^59^ Subsequently, xAM images, additional B-mode images, and finally BURST images were acquired at the same location without moving the transducer using the same parameters as described above for the *in vitro* experiments (e.g. 18V for xAM, 1.6V for B-mode, and 1.6V to 25V for BURST). At least two separate locations spaced at least 2 mm apart in each tumor were imaged with B-mode, pAM, and xAM. Ultrasound images of tumors were quantified as described above where the sample ROIs were drawn around the necrotic cores in the tumors and the background ROIs were drawn around regions in the gel above the mouse. Images were normalized and plotted on a dB scale as described above except the scaling factors were A=2.5 and B=1 for xAM and pAM and the corresponding B-mode tumor images, and A=10 and B=0.5 for BURST images.

### Histology of tumors colonized by bacteria

Tumors were colonized with pBAD-bARG_Ser_-AxeTxe EcN following the same protocol as described above. The day after inducing GV expression with IP injections of L-arabinose, BURST, xAM and B-mode images of tumors were acquired as described above. Shortly after imaging, mice were euthanized by sedation with isoflurane and cervical dislocation. Tumors were resected, placed in 10% buffered formalin for 48 hours, and then washed and stored in 70% ethanol. Tumors were then cut in half along the approximate plane of imaging, placed in tissue cassettes, and sent to the Translational Pathology Core Laboratory at UCLA, which embedded samples in paraffin and performed H&E staining, immunohistochemistry, and microscopy imaging. Immunohistochemistry was performed using Opal IHC kits (Akoya Biosciences) according to the manufacturer’s instructions. Tissue sections were incubated with either polyclonal rabbit anti-*E. coli* antibody (Virostat; catalogue number 1001) or non-reactive rabbit IgG isotype control antibody as a negative control. All sections were then incubated with an Opal 520 polymer anti-rabbit HRP antibody (Akoya Biosciences) and counterstained with DAPI. Sections were imaged in the appropriate fluorescence or brightfield channels using a high throughput scanning system (Leica Aperio VERSA) with 40 μm resolution.

### Mammalian plasmid construction

Monocistronic plasmids were constructed using standard cloning techniques including Gibson assembly and conventional restriction and ligation. Coding sequences for the *A. flos-aquae* GV genes were codon-optimized and synthesized by Integrated DNA Technologies, and subcloned into a pCMVSport backbone with a CMV promoter as described previously.^7^ *gvpA-*WPRE-hGH polyA was constructed by subcloning *gvpA* between PstI and MluI sites of pCMVSport vector with WPRE-hGH polyA.

Polycistronic mARG_Ana_ assembly factor genes *gvpNJKFGWV* were synthesized by Twist Bioscience in a pTwist-CMV vector. Emerald GFP was subcloned in-frame downstream of the *gvpNJKFGWV* ORF via a P2A linker and the entire *pNJKFGWV-GFP* ORF was subcloned into a pCMVSport backbone with WPRE-hGH-poly(A) elements using NEBuilder HiFi DNA Assembly (NEB). *gvpA-*IRES-EBFP2-WPRE-hGH polyA was constructed by Gibson assembly of a PCR-amplified IRES-EBFP2 fragment into the XbaI site of *gvpA--*WPRE-hGH polyA plasmid.

PiggyBac transposon plasmids were constructed by PCR amplifying the region between the start codon of *gvpNJKFGWV*or *gvpA* and the end of the hGH poly(A) from the pCMVSport plasmids. The amplified regions were Gibson-assembled into the PiggyBac transposon backbone (System Biosciences) with a TRE3G promoter (Takara Bio) for doxycycline-inducible expression.

The lentiviral transfer plasmid with constitutively expressed tetracycline transactivator (pEF1α-rtTA-Antares-WPRE) was constructed as follows: pNCS-Antares was obtained from Addgene (#74279) and P2A was added to the N-terminus of Antares with a primer overhang during PCR. This fragment was subcloned into the lentiviral transfer plasmid pEF1α-rtTA-WPRE between rtTA and WPRE inframe with the rtTA ORF using NEBuilder HiFi DNA Assembly.

### HEK293T cell culture, transient transfection, in vitro ultrasound imaging of transient expression of GVs suspended in agarose phantoms

HEK293T cells (ATCC, CLR-2316) were cultured in 24-well plates at 37°C, 5% CO2 in a humidified incubator in 0.5ml DMEM (Corning, 10-013-CV) with 10% FBS (Gibco) and 1x penicillin/streptomycin until about 80% confluency before transfection as described previously.^7^ Briefly, transient transfection mixtures were created by mixing of around 600 ng of plasmid mixture with polyethyleneimine (PEI-MAX; Polysciences Inc.) at 2.58 μg polyethyleneimine per μg of DNA. The mixture was incubated for 12 minutes at room temperature and added drop-wise to HEK293T cells. Media was changed after 12-16 hours and daily thereafter. For *gvpA* titration experiments, pUC19 plasmid DNA was used to keep the total amount of DNA constant.

After three days of expression, cells were dissociated using Trypsin/EDTA, counted using disposable hemocytometers (Bulldog), and centrifuged at 300g for 6 minutes at room temperature. Cells were resuspended with 1% low-melt agarose (GoldBio) in PBS at 40°C at ~30 million cells/mL (**Fig. 5b,f**,~15 million cells/mL (**Fig. 5d**), or ~7.5 million cells/mL (Fig S18c) before loading into wells of preformed phantoms consisting of 1% agarose (Bio-Rad) in PBS.

Phantoms were imaged using L22-14v transducer (Verasonics) while submerged in PBS on top of an acoustic absorber pad. For BURST imaging, wells were centered around the 8 mm natural focus of the transducer and a BURST pulse sequence was applied in pAM acquisition mode as described above, except the focus was set to 8 mm, and the acoustic pressure was set to 0.26 MPa (1.6V) for the first 10 frames and 2.11 MPa (10V) for the remaining frames. **The** xAM voltage ramps and B-mode images were acquired concurrently using the same parameters as described above, except the transducer voltage was varied from 4 to 24V in steps of 0.5V for xAM, and 10 frames, each consisting of 15 accumulations, were acquired per voltage. The well depth and the B-mode transmit focus were set to 5 mm. All image quantification was performed as described above, where the sample ROIs were drawn inside the well and the background ROIs were drawn around an empty region in the agarose phantom for SBR calculation. All images were normalized and plotted on a dB scale as described above except the scaling factors were A=2 and B=0.5.

### Genomic integration and FACS

MDA-MB-231 (ATCC, HTB26), 3T3 (ATCC, CRL-1658) and HuH7 (JCRB0403) cells were cultured in DMEM (Corning, 10-013-CV) supplemented with 10% FBS (Gibco) and 1x penicillin/streptomycin at 37°C and 5% CO2 in a humidified incubator unless noted otherwise. Cells were lentivirally transduced with pEF1α-rtTA-Antares-WPRE and sorted based on strong Antares fluorescence (Ex: 488 nm, Em: 610/20BP + 595LP) using BD FACSAria II for MDA-MB-231 cells and MACSQuant Tyto (B2 channel) for 3T3 and HuH7 cells. MDA-MB-231-rtTA-Antares cells were then electroporated in 20ul format using 4D-Nucleofector using CH-125 protocol in SF buffer (Lonza) with 1 μg PiggyBac transposon:transposase plasmid mixture (2:1 PB-gvpA:PB-gvpNV transposons, 285ng PiggyBac transposase). 3T3-rtTA-Antares cells were transfected with the same PiggyBac plasmid mixture using PEI-MAX in a 12-well format and HuH7 cells were transfected using Lipofectamine 3000. Cells were expanded into surface-treated T75 flasks in TET-free media and were induced for 12 hours with 1 μg/mL doxycycline before sorting for triple positive cells (gated for Antares, then Emerald and EBFP2). The sorted cells were returned to DMEM with TET-free FBS (Takara). MDA-MB-231-mARG_Ana_ was sorted twice. The first round of sorting was performed with permissive gates and the enriched population was ~50% double positive for Emerald and EBFP2 as analyzed with MACSQuant VYB (Miltenyi Biotec). This population was sorted again with stricter gates to ~95% purity. 3T3-mARG_Ana_ cells were sorted for strong Emerald and EBFP2 fluorescence using MACSQuant Tyto only once, yielding a population that was ~80% double positive. HuH7 cells were sorted twice to ~91% purity using MACSQuant Tyto. Cells were expanded in TET-free media and frozen in Recovery Cell Culture Freezing Medium (Gibco) using Mr. Frosty cell freezing container (Nalgene) filled with isopropanol at −80°C, and then stored in liquid nitrogen vapor phase until use.

### In vitro ultrasound imaging of MDA-MB-231 mARG_Ana_ cells suspended in agarose phantoms

For all *in vitro* experiments, MDA-MB-231-mARG_Ana_ cells were cultured in DMEM supplemented with 10% TET-free FBS and penicillin/streptomycin. For xAM imaging of MDA-MB-231-mARG_Ana_ cells suspended in agarose phantoms, cells were cultured in 24-well plates in 0.5 mL media. For **Fig. 5h**, cells were seeded at 7,500 cells per well and induced with 1 μg/mL doxycycline after an overnight incubation and at subsequent days as indicated (5 replicates per condition), except for the uninduced control which was grown in a 10 cm dish without doxycycline. Media was changed daily thereafter until cell harvest. Cells were trypsinized with 100 μL Trypsin/EDTA for 6 minutes at 37°C, after which the trypsin was quenched by addition of 900 μL media. The cell number was equalized between different days of expression at 140,000 cells and pelleted at 300g for 6 minutes. Cells were then resuspended in 20 μl 1% low-melt agarose (GoldBio) in PBS at 40°C and loaded into the wells of preformed 1% agarose (Bio-Rad) phantoms in PBS. Ultrasound images were acquired with L22-14v 128-element linear array transducer (Verasonics). xAM voltage ramp and B-mode images were acquired concurrently using the same parameters as described above (the transducer voltage was varied from 4 to 24V in steps of 0.5V for xAM and 10 frames, each consisting of 15 accumulations, were acquired per voltage. The B-mode transmit focus was set to 5 mm). Images taken at the voltage that produced peak xAM signal (9V, 0.54 MPa) were chosen for quantification. For **Fig. 5i,j**, cells were seeded at 66,666 cells per well and induced with the indicated doxycycline concentrations after an overnight incubation in TET-free media (4 replicates per doxycycline concentration). Cells were incubated for 4 days with daily media/doxycycline changes. Cells were harvested as above, and ~420,000 cells from each condition were loaded per agarose phantom well. xAM and B-mode images were acquired concurrently using the same parameters as described above except the transducer voltage was varied from 6V to 10V in steps of 0.5 V for xAM and 120 frames, each consisting of 15 accumulations, were acquired per voltage (~75 seconds/voltage). The B-mode transmit focus was set to 6 mm. Images taken at 7.5V (0.42 MPa) were chosen for display and quantification in **Fig. 5i** (doxycycline response). For **Fig. 5k** and **Fig. S18f-g,** cells were seeded in 10-cm dishes and induced as above for 4 days. Cells were harvested as above and resuspended at 60,000,000 cells/mL. 10-fold serial dilutions were performed with each cell line. Each cell dilution was mixed 1:1 with 2% low-melt agarose before loading into agarose phantom wells. Cells were imaged with an L22-14vX transducer at 5.5V (0.61 MPa) for xAM: the highest pressure that produced stable signal over a 30-second exposure and using 2V to 15V pAM BURST.

For imaging of MDA-MB-231 cells under thick liver tissue, cells were induced with doxycycline in T225 flasks for 4 days. Cells were harvested as above and resuspended at 30,000,000 cells/mL in 1% low-melt agarose in PBS prior to loading into agarose phantom wells. > 1 cm beef liver section (99 Ranch Market) was overlaid on top of the agarose phantom and secured with needles. The phantom and liver were submerged in a PBS bath and the transducer was positioned 20 mm away from the interface between the liver and the agarose phantom. Ultrasound imaging was performed using a L10-4v linear array transducer (Verasonics) using the same parameters as above, except the xAM voltage was varied between 2V (0.078 MPa) and 30V (2.51 MPa). B-mode was acquired at 1.6V (0.25 MPa). Each voltage was held for 5 frames, each consisting of 15 accumulations.

All image quantification was performed as described above where the sample ROIs were drawn inside the well and the background ROIs were drawn around an empty region in the agarose phantom for SBR calculation. All Images were normalized and plotted on a dB scale as described above except the scaling factors were A=2 and B=0.5. The xAM/B-mode overlay was made with the B-mode image as background. A binary alpha mask was applied to the xAM image, giving pixel values lower than 2x the average background a value of 0 and all values above this threshold a value of 1.

### TEM imaging of GVs expressed in mammalian cells

For TEM, cells were cultured in 6-well plates in 2 mL media. 1 μg/mL doxycycline was added to the wells at indicated times with daily media plus doxycycline changes thereafter until harvest. Cells were lysed by adding 400 μL of Solulyse-M (Genlantis) supplemented with 25 units/mL Benzonase Nuclease (Novagen) directly to the 6-well plates and incubating for 1 hour at 4°C with agitation. The lysates were then transferred to 1.5 mL microcentrifuge tubes. Eight hundred μL of 10 mM HEPES pH 7.5 was added to each tube, and lysates were centrifuged overnight at 300g and 8°C. Thirty μL of the supernatant was collected from the surface from the side of the tube facing the center of the centrifuge rotor and transferred to a new tube. Three μL of each sample was loaded onto freshly glow-discharged (Pelco EasiGlow, 15mA, 1 min) formvar/carbon 200 mesh grids (Ted Pella) and blotted after 1 minute then air-dried. The unstained grids were imaged on a FEI Tecnai T12 transmission electron microscope equipped with a Gatan Ultrascan CCD.

### In vivo ultrasound imaging of mARG_Ana_ expressing orthotopic tumors, whole animal fluorescence imaging and tumor fluorescence microscopy

Tumor xenograft experiments were conducted in NSG mice aged 12-weeks and 6 days (The Jackson Laboratory). To implement an orthotopic model of breast cancer, all the mice were female. MDA-MB-231-mARG_Ana_ cells were grown in T225 flasks in DMEM supplemented with 10% TET-free FBS and penicillin/streptomycin until confluency as described above. Cells were harvested by trypsinization with 6 mL Trypsin/EDTA for 6 minutes and quenched with fresh media. Cells were washed once in DMEM without antibiotics or FBS before pelleting by centrifugation at 300g. Cell pellets were resuspended in 1:1 mixture of ice-cold Matrigel (HC, GFR) (Corning 354263) and PBS (Ca^2+^, Mg^2+^-free) at 30 million cells/mL. Fifty μL Matrigel suspensions were injected bilaterally into the 4^th^ mammary fat pads at 1.5 million cells per tumor via subcutaneous injection. 12 hours after tumor injection and every 12 hours thereafter (except the mornings of ultrasound imaging sessions) test mice were intraperitoneally injected with 150 μL of saline containing 150 μg of doxycycline for induction of GV expression. Control mice were not injected with doxycycline.

For ultrasound imaging, mice were depilated around the 4^th^ mammary fat pads using Nair (Aloe Vera) for ultrasound coupling with Aquasonic 100 gel. Mice were anesthetized with 2.5% isoflurane and maintained at 37°C in supine position on a heating pad. The first imaging session (day 4) consisted of 8 induced tumors from 4 mice and 7 uninduced tumors from 4 mice. One of the uninduced mice died during the first imaging session, which resulted in two fewer uninduced control tumors for the remaining imaging sessions.

Ultrasound images were acquired with an L22-14v 128-element linear array transducer. xAM and B-mode images were acquired concurrently using the same parameters as described in the *in vitro* section above except the transducer voltage was held at constant 7.5V (0.42 MPa) for xAM and 3 frames, each consisting of 15 accumulations, were acquired per section. A motor stage was programed to move 100 μm per section for a total of 150 sections per tumor. The B-mode transmit focus was set to 6 mm. Ultrasound images of tumors were quantified as described above where the sample ROIs were drawn around the tumors and the background ROIs were drawn around regions in the gel above the mouse. Images were normalized and plotted on a dB scale as described above except the scaling factors were A=2 and B=0.5 for both xAM and the corresponding B-mode tumor images. The xAM volume quantification was performed by summing all pixel values from all sections in each tomogram between 2 mm and 10 mm in depth.

On the last day of ultrasound imaging, mice were anesthetized with 100 mg/kg ketamine, and 10 mg/kg xylazine and whole-body imaged in supine position using ChemiDoc MP imaging system with Image Lab software (BIO-RAD). Fluorescence channels were set as follows: blue epi illumination with 530/28 filter for Emerald/GFP and 605/50 filter for Antares/CyOFP1. Images were processed and merged using the FIJI package of ImageJ.

After whole-body fluorescence imaging, mice were euthanized and tumors were resected and placed in 10% formalin solution for 24 hours at 4°C, after which they were transferred to PBS. Fixed tumors were embedded in 2% agarose in PBS and sectioned to 100 μm slices using a vibratome. Sections were stained with TO-PRO-3 nucleus stain, mounted using Prolong Glass (Invitrogen) and imaged using a Zeiss LSM 980 confocal microscope with ZEN Blue. Images were processed using the FIJI package of ImageJ. For micrographs of tumors from both induced and uninduced mice, the Emerald channel was capped between 0 and 25497, EBFP2 channel between 0 and 17233 and TO-PRO-3 channel between 5945 and 53136 for display.

### In vivo *ultrasound-guided biopsy of mARG_Ana_-expressing chimeric tumors*

Chimeric tumor biopsy experiments were conducted in female NCG mice aged 8-weeks (Charles River Laboratories). MDA-MB-231-mARG_Ana_ and MDA-MB-231-rtTA-Antares cells were grown and harvested as above. Cell pellets were resuspended in a 1:1 mixture of ice-cold Matrigel (HC, GFR) (Corning 354263) and PBS (Ca^2+^- and Mg^2+^-free) at 30 million cells/mL. 100 μL Matrigel suspensions of MDA-MB-231-mARG_Ana_ were injected bilaterally into the 4^th^ mammary fat pads at 3 million cells per tumor lobe via subcutaneous injection. After 1 hr, additional 100 μL Matrigel suspensions of MDA-MB-231-rtTA-Antares were injected close to the edge of the blisters created by the first injections to create dual-lobed chimeric tumors with heterogeneous gene expression patterns. Mice were intraperitoneally injected with 150 μL of saline containing 150 μg of doxycycline for induction of GV expression starting 12 hours after tumor injection and then every 12 hours thereafter for 5 days.

Mice were prepared for ultrasound imaging as above. Ultrasound images were acquired with an L22-14vX 128-element linear array transducer. xAM and B-mode imaging was performed as above except the transducer voltage was held at 5.5V (0.481 MPa) for xAM and 1.6V (0.161 MPa) for B-mode and the motor stage was programed to move either 200 μm per section for whole-tumor scans or was held stationary for biopsy video acquisition. Image normalization and scaling was performed as above. xAM/B-mode overlay was made as above.

To perform a fine-needle aspiration biopsy, a 23G needle was fitted to a 3 mL Luer-lock syringe prefilled with PBS. The syringe was mounted on a 3D-printed holder attached to a manual translation stage. Each biopsy attempt consisted of positioning the ultrasound probe over a tumor and moving the needle into the field of view. The needle was then inserted into either the xAM-positive or xAM-negative region of the tumor, guided by live xAM and B-mode imaging. The needle was wiggled back-and-forth a couple times before pulling the syringe plunger to aspirate cells. The tumor sample was then ejected into a tube with PBS. The biopsy was repeated for attempts that did not produce a visible cell pellet. Each sample was treated with Trypsin/EDTA for 6 minutes at 37°C, then quenched with fresh media.

Flow cytometry was performed with MACSQuant 10 (Miltenyi Biotec). GFP was measured with the B1 channel and Antares using B2. All biopsy attempts for a given tumor/sampling condition were analyzed separately, but their resulting FCS data files were concatenated. Data analysis was performed in FlowJo. For quantification of biopsy samples, each population was first gated for Antares-positive cells to exclude endogenous mouse cells. Antares-positive cells were then gated based on FSC/SSC and single cells were gated using FSC-A vs FSC-W. The resulting populations contained on average 6947 cells with a SD of 6960 cells and range between 72 and 21958 cells. %GFP-positive (mARG_Ana_-positive) was assessed based on these resulting populations.

## Data availability

Plasmids will be made available through Addgene upon publication. All other materials and data are available from the corresponding author upon reasonable request.

## Code availability

Ultrasound data acquisition and analysis code will be made available on the Shapiro Lab GitHub at https://github.com/shapiro-lab upon publication.

## ACKNOWLEDGEMENTS

The authors would like to thank Dianne Newman for a sample of *Streptomyces coelicolor* A3(2), and Yunfeng Li, Avinoam Bar-Zion and Hongyi (Richard) Li for help with tissue histology. Electron microscopy was performed in the Beckman Institute Resource Center for Transmission Electron Microscopy at Caltech. Mammalian cell sorting was performed at the Analytical Cytometry Core at City of Hope in Duarte, CA. Confocal microscopy was performed in the Beckman Institute Biological Imaging Center. This research was supported by the National Institutes of Health (R01-EB018975 to M.G.S.) and Pew Charitable Trust. R.C.H. was supported by the Caltech Center for Environmental Microbial Interactions. M.T.B. was supported by an NSF GRFP fellowship. Related research in the Shapiro Laboratory is supported by the David and Lucille Packard Foundation, the Burroughs Wellcome Fund and the Heritage Medical Research Institute.

## AUTHOR CONTRIBUTIONS

R.C.H., M.T.B., M.D., K.W., M.B.S., P.D., Z.J., M.Y.Y., A.F. and R.D. planned and performed experiments. R.C.H. conceived and performed the phylogenetic screening experiments. M.T.B. and M.D. performed all *in vivo* experiments, with help from M.B.S and P.B.-L.. P.D. and M.D. performed TEM imaging. D.R.M. built the ultrasound plate-scanning setup, and D.R.M. and D.P.S. wrote the associated MATLAB scripts for controlling it. Z.J. and D.P.S. wrote the MATLAB scripts for ultrasound imaging of EcN *in vitro* and *in vivo*. Z.J. performed the calibration of the L22-14v transducer. M.H.A. provided the axe-txe stability cassette, and advised on tumor colonization experiments. R.C.H., M.T.B., M.D., D.P.S., P.D., and Z.J. analyzed data. R.C.H., M.T.B., M.D., and M.G.S. wrote the manuscript with input from all other authors. M.G.S. supervised the research.

## COMPETING INTERESTS

The authors declare no competing financial interests.

## SUPPLEMENTARY INFORMATION

Supplementary Figures 1-20 Supplementary Table 1

Supplementary Video 1 (xAM/B-mode tomogram of induced orthotopic tumor on day 12)

Supplementary Video 2 (xAM/B-mode tomogram of induced chimeric tumor on day 5)

Supplementary Video 3 (xAM/B-mode 3D reconstruction of the tumor from Video 2)

Supplementary Video 4 (xAM/B-mode video of chimeric tumor biopsy procedure sampling of xAM-positive region)

Supplementary Video 5 (xAM/B-mode video of chimeric tumor biopsy procedure sampling of xAM-negative region)

## SUPPLEMENTARY INFORMATION

**Figure S1:**
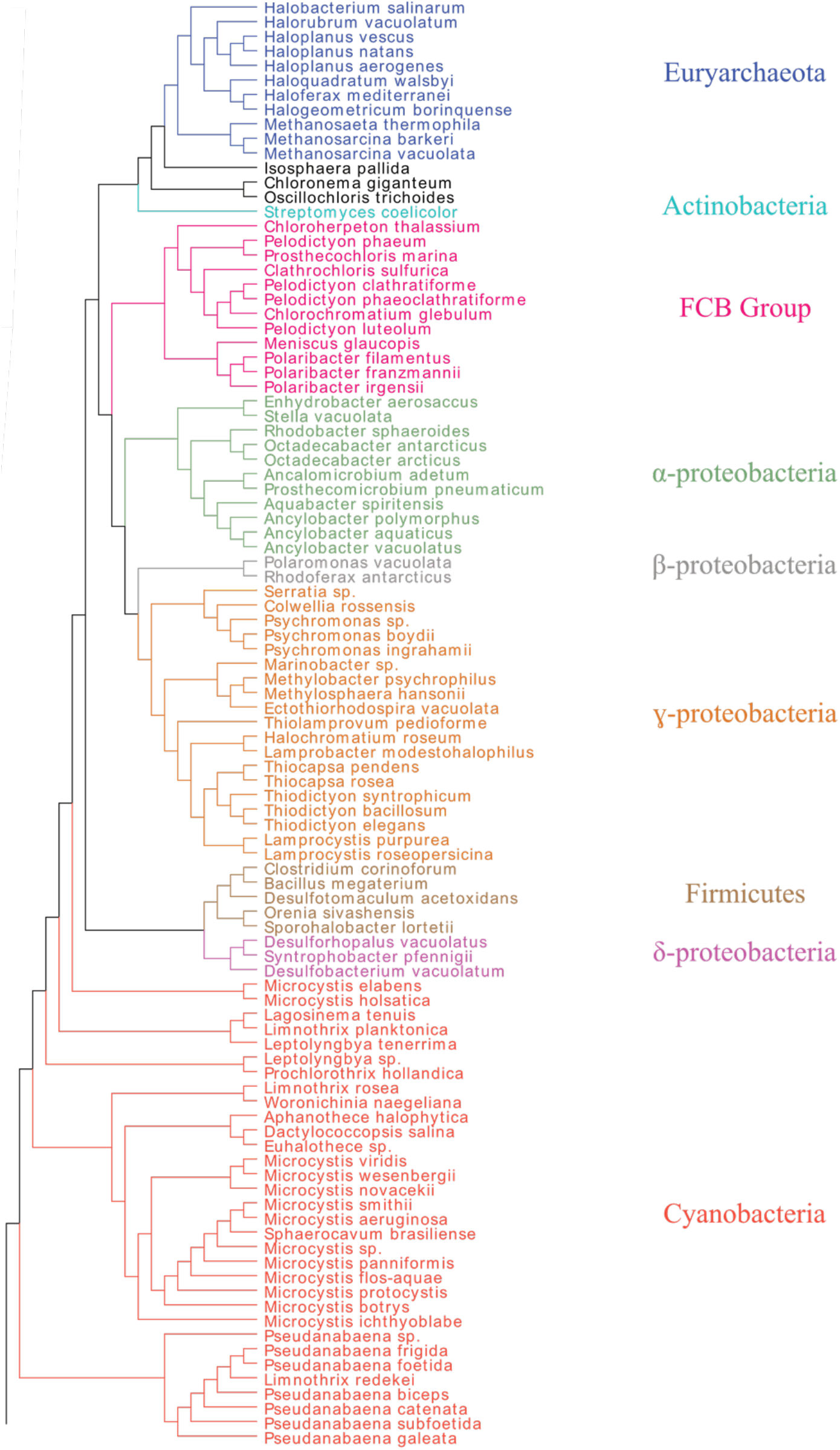
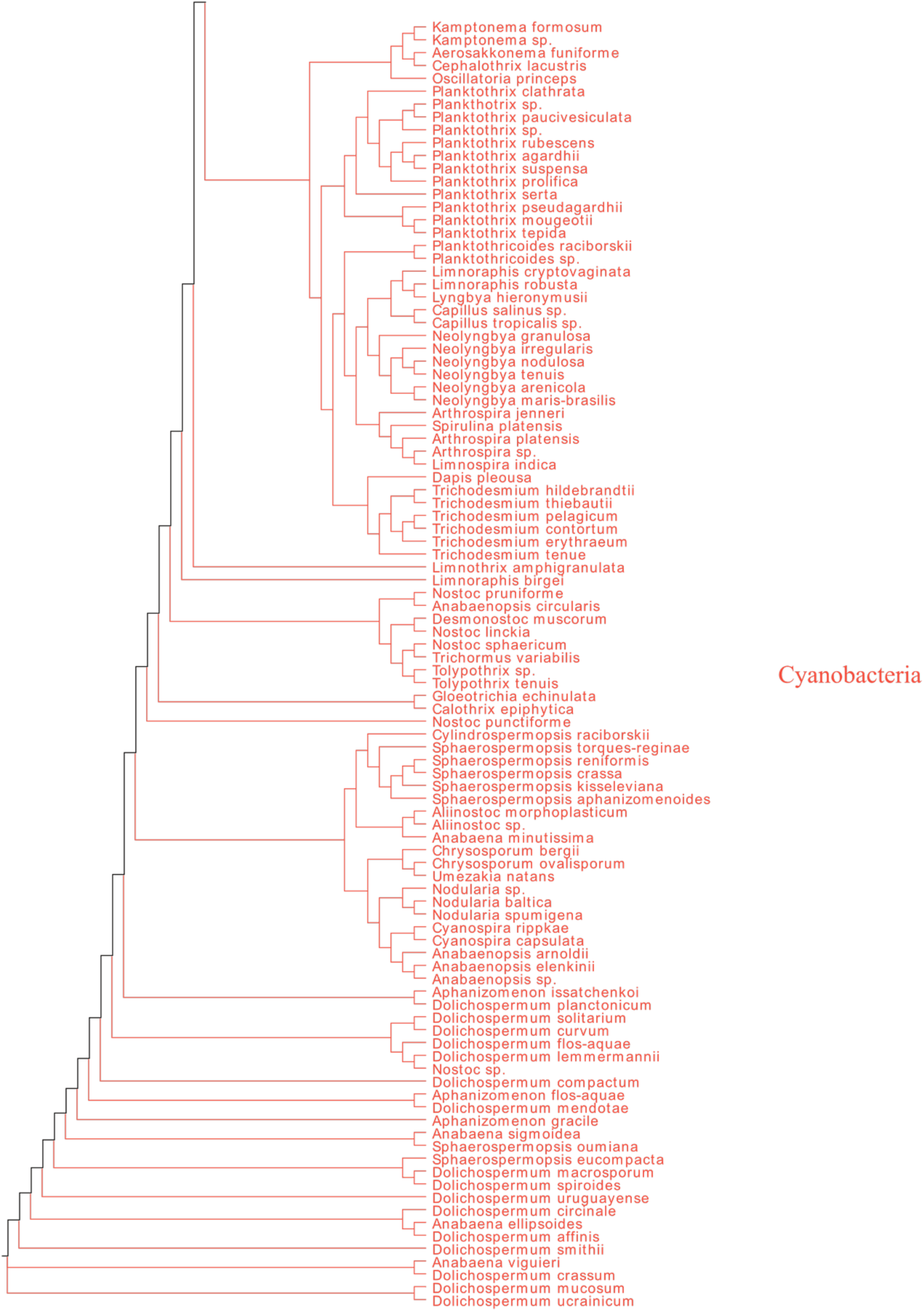
16S phylogenetic tree of all reported GV-producing organisms. Colors indicate groupings of phylogenetically similar organisms. Organisms from which GV genes were tested in *E. coli* are shown in **Fig. 1a**.

**Figure S2:**
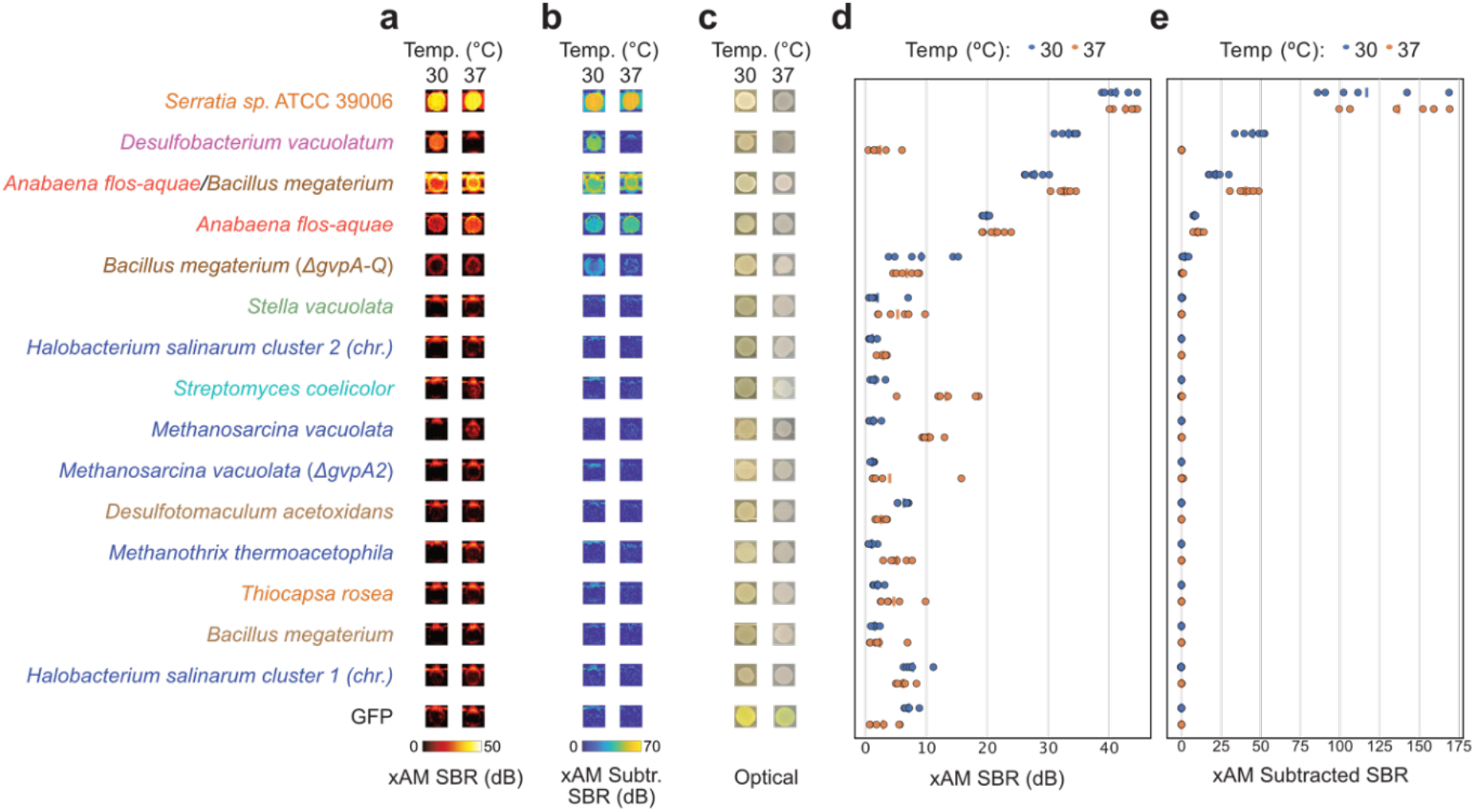
Additional images and quantification of *E. coli* patches expressing select GV gene clusters. **(a-c)** xAM images (a), pre-minus-post-collapse xAM images (b), and optical images showing opacity (c) of patches of *E. coli* expressing various GV gene clusters from the organisms listed on the left. (**de**) Quantification of the xAM (d) and pre-minus-post-collapse xAM (e) signals from images in (a-b) (n=6). SBR, signal-to-background ratio.

**Figure S3:**
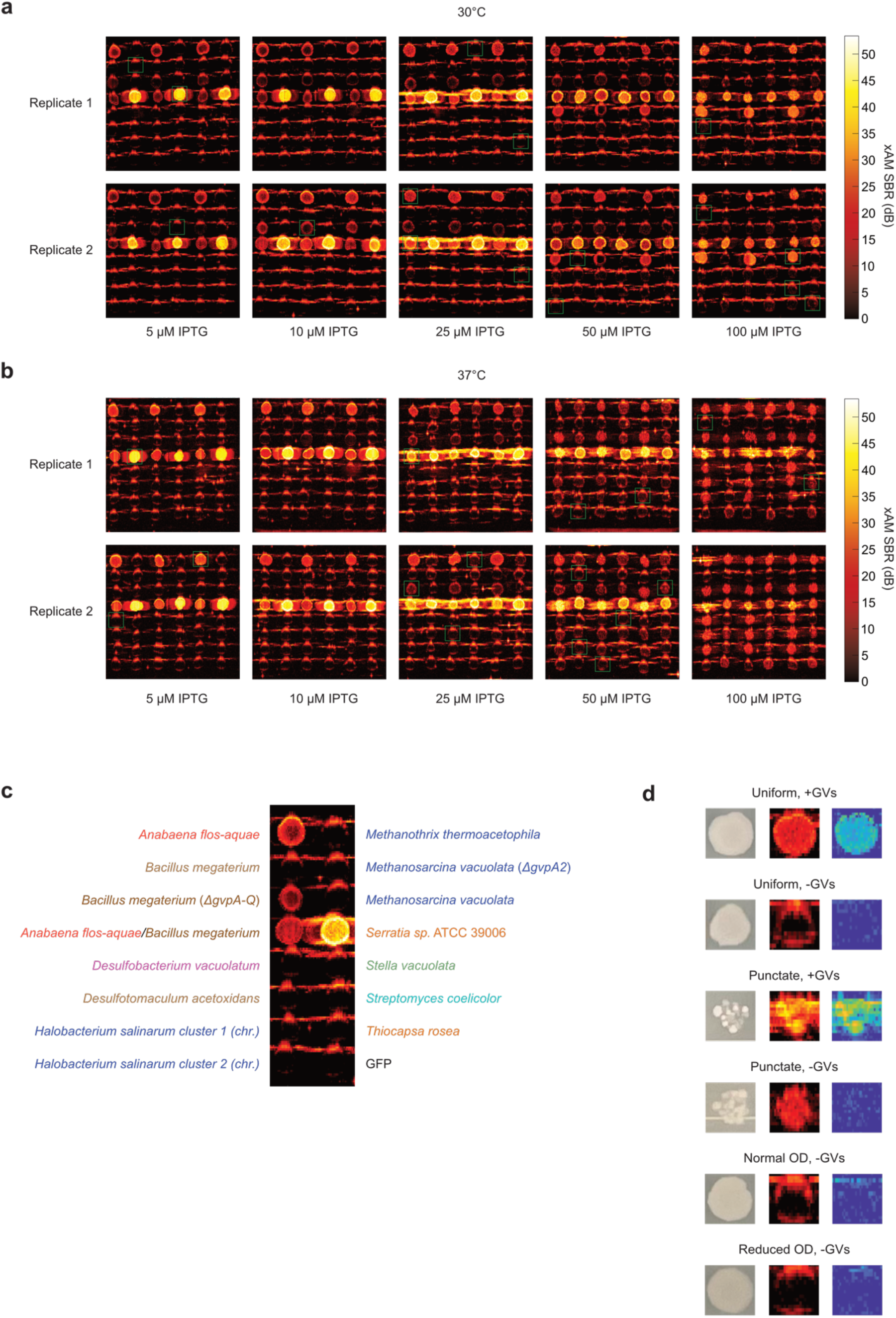
Optimization of expression conditions for all tested clusters in BL21(DE3) *E. coli.* (**a-b**) xAM images of bacterial patches expressing each GV cluster at varying inducer concentrations and temperatures. Green boxes indicate the patches shown in **Fig. S2a**. The IPTG concentration selected was the one that resulted in the highest xAM pre-minus-post-collapse difference signal (**Fig. S6**) while not creating toxicity, as determined by whether the patch was uniform or punctate (**Fig. S5a-b**). Some of the IPTG concentrations that led to toxicity also created significant xAM signal, but this signal did not originate from GVs, as indicated by the lack of xAM pre-minus-post-collapse signal difference (**Fig. S6**). Further, there were some IPTG concentrations for certain genotypes that created significant xAM signal but no xAM pre-minus-post-collapse signal difference, and no visible toxicity (e.g., *Streptomyces coelicolor, Thiocapsa rosea,* and GFP at 37°C, 100 μM IPTG). This discrepancy was likely caused by subtle toxicity that is not apparent in optical images, but altered the texture of the patch enough to be detectable by US. (**c**) Key for genotypes tested in (a-b), with this pattern repeated in three pairs of columns replicated on each plate. (**d**) Examples of the effects of toxic genotypes on bacterial patches, and of artifacts that can appear in bacterial patch scan images. Bacteria themselves can produce significant xAM signal (especially when present in extremely high concentrations, as they are in the confluent patches imaged here), which can be seen in the forms of rings around all patches, regardless of GV expression status. Further, expression of toxic proteins (or of large amounts of otherwise non-toxic proteins, such as GFP) can interfere with bacterial growth; in extreme cases this results in significant cell death and a punctate appearance, and in less extreme cases it simply reduces the optical density of patches. GV expression can increase the optical density of patches, but only at high levels of GV expression. Punctate patches produce considerably more xAM signal than uniform ones, even in the absence of GV expression. The xAM pre-minus-post-collapse difference can be used to qualitatively determine if a patch produces GVs, but because collapse is incomplete in some cases, it is not an ideal method for quantitatively comparing genotypes.

**Figure S4:**
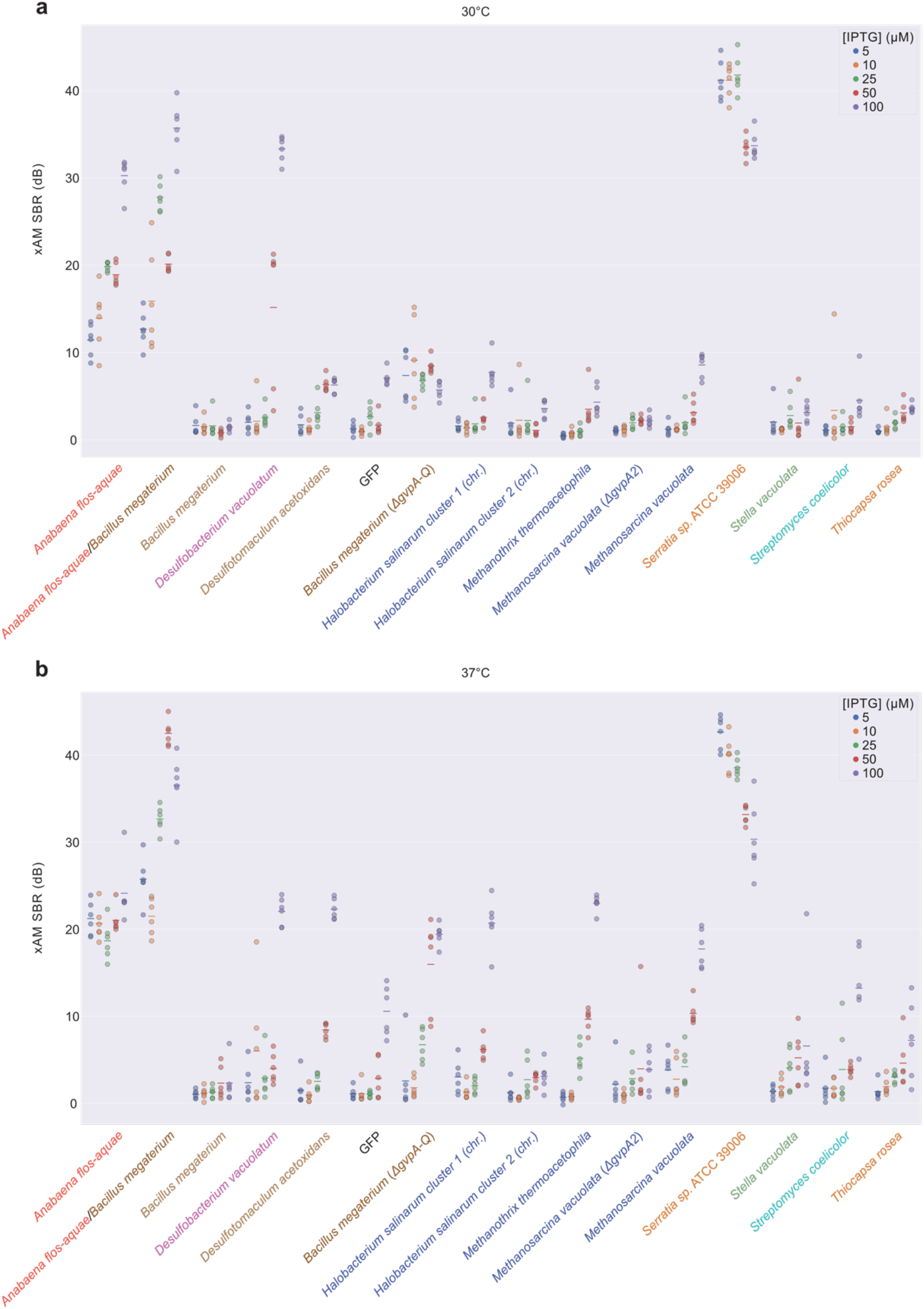
Quantification of ultrasound signal for all samples shown in Fig. S3a-b. (**a-b**) xAM SBR of the patches at 30°C (a) and 37°C (b) shown in **Fig. S3a-b** (n=6; lines represent the mean).

**Figure S5:**
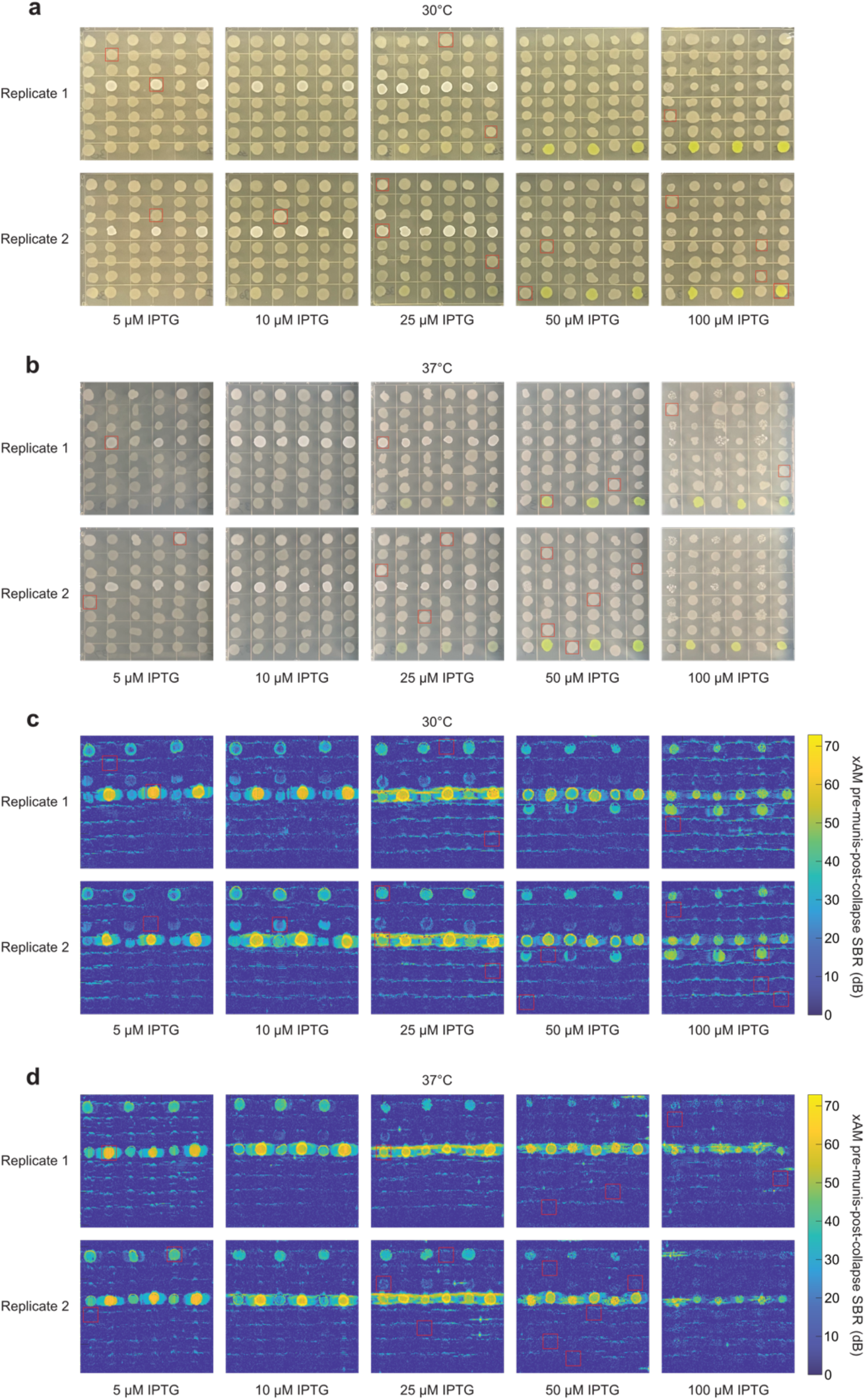
Optical and xAM pre-minus-post-collapse difference images of all samples shown in Fig. S3. (**a-b**) Optical images of patches at 30°C (a) and 37°C (b) shown in **Fig. S3a-b**. (**c-d**) xAM pre-minus-post-collapse difference patches of samples at 30°C (a) and 37°C (b) shown in **Fig. S3a-b**. Red boxes indicate the patches shown in **Fig. S2b-c**.

**Figure S6:**
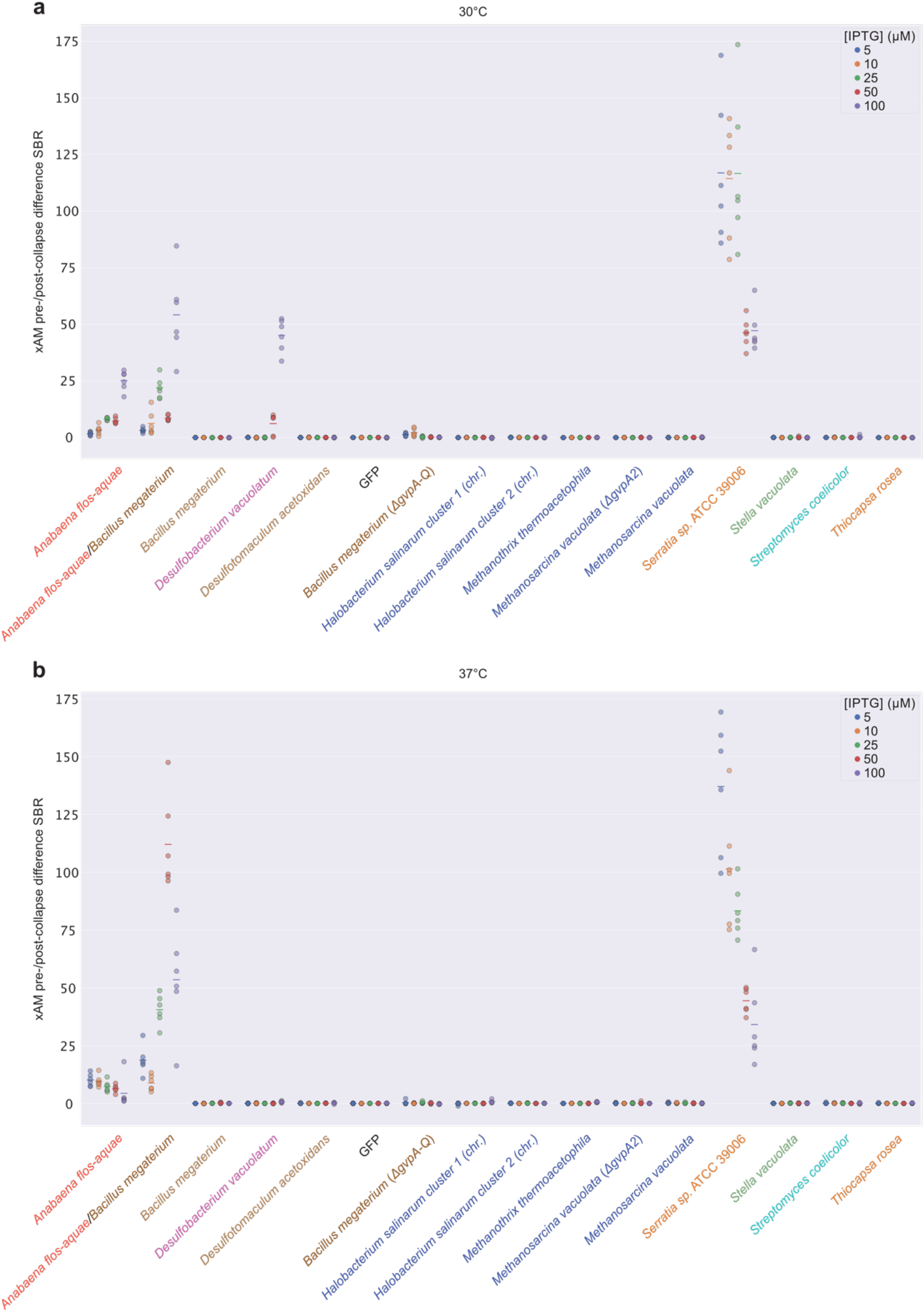
Quantification of ultrasound signal for samples shown in Fig. S5c-d. (**a-b**) xAM SBR for the patches at 30°C (a) and 37°C (b) shown in **Fig. S5c-d** (n=6; lines represent the mean).

**Figure S7:**
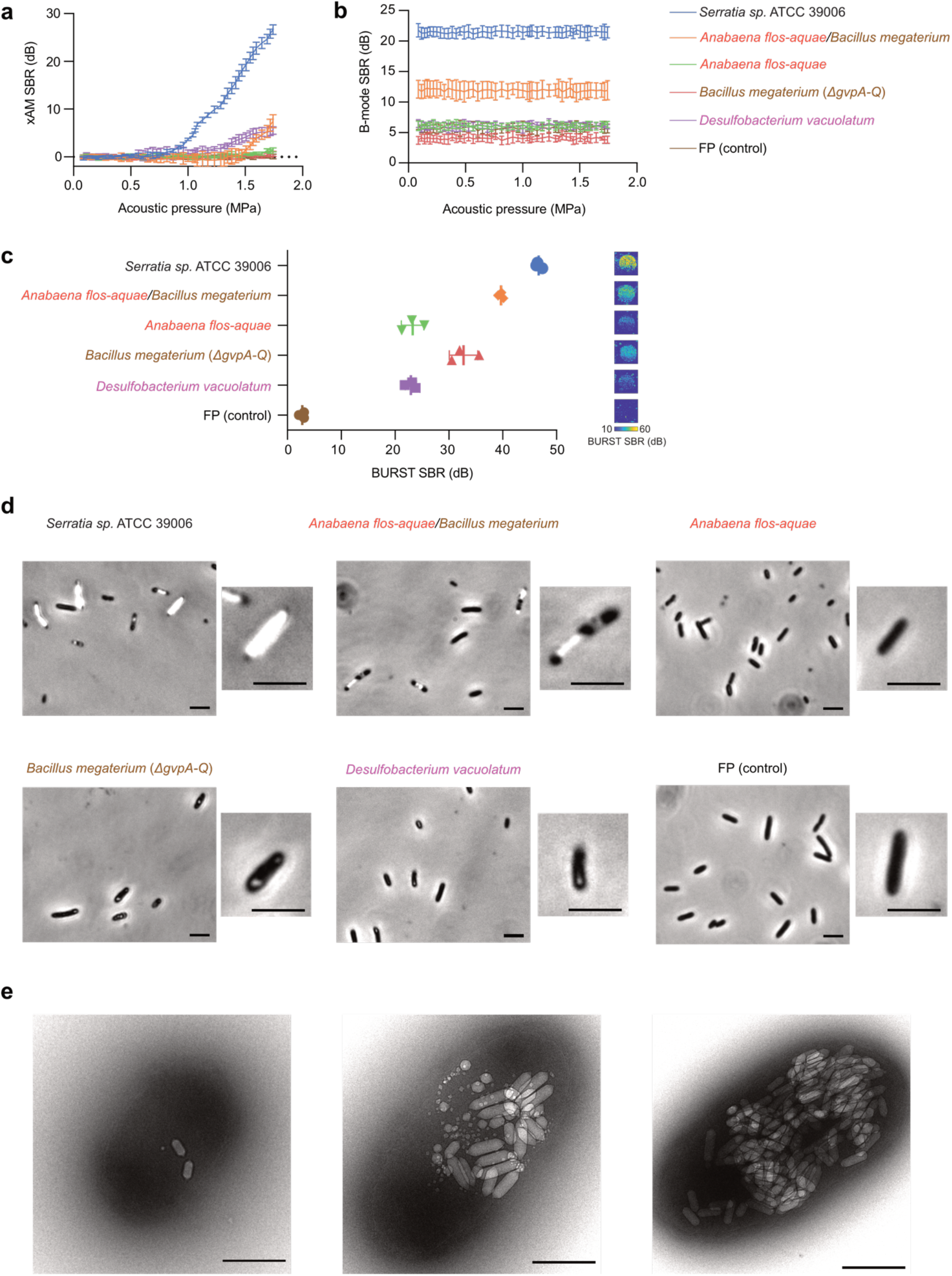
Characterization of working GV clusters in BL21(DE3) *E. coli*. (**a-d**) xAM signal-to-background ratio (SBR) as a function of acoustic pressure (a), B-mode SBR at a constant pressure of 0.15 MPa after each increase in acoustic pressure in a (b), BURST SBRs and corresponding representative images (c), and representative phase contrast microscopy (PCM) images (d) of the working GV clusters expressed in BL21(DE3) *E. coli* at 30°C on solid media. For ultrasound imaging (a-c), samples were normalized to 5 × 10^9^ cells/mL in agarose phantoms. Curves and error bars represent the mean (n=3 biological replicates each with 2 technical replicates) ± SD. (a-b) have the same legend. GV clusters in cells are visible by PCM for all clusters except for the cluster from *Anabaena flos-aquae* and the fluorescent protein (FP) control (d). (**e**) Representative TEM images of BL21(DE3) *E. coli* cells expressing Serratia GVs at varying levels of expression. Scale bars are 5 μm in (d) and 500 nm in (e).

**Figure S8:**
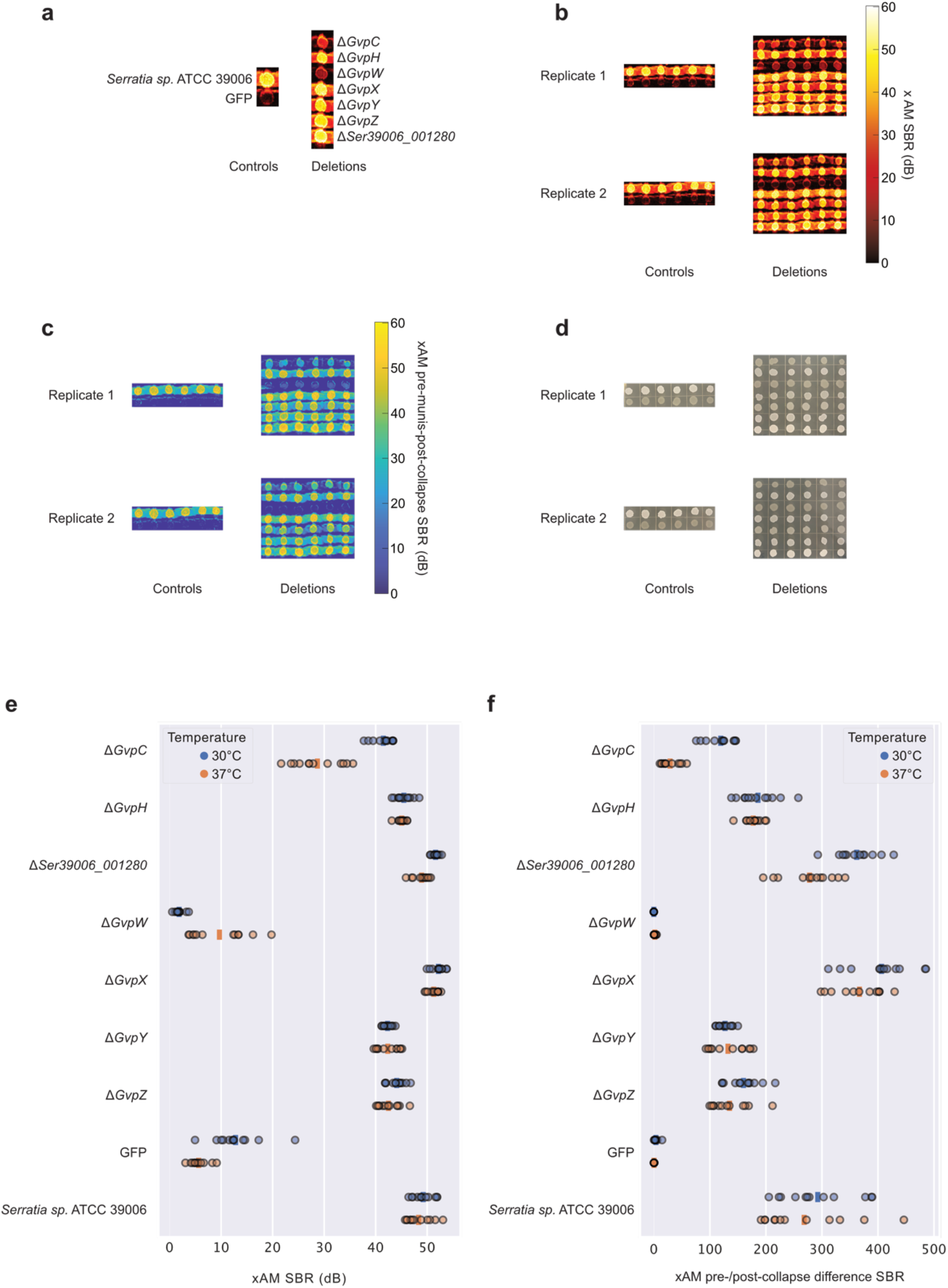
Effects of single-gene deletions on GV expression by the *Serratia* cluster. (**a**) Key for genotypes tested, repeated in 6 replicate columns on each plate. (**b-d**) xAM images (b), pre-minus-post-collapse xAM images (c), and optical images (d) of bacterial patches expressing single-gene deletions of the Serratia cluster. (**e-f** Quantification of the xAM images (e) and pre-minus-post-collapse xAM images (f) shown in (b-c) (n=12).

**Figure S9:**
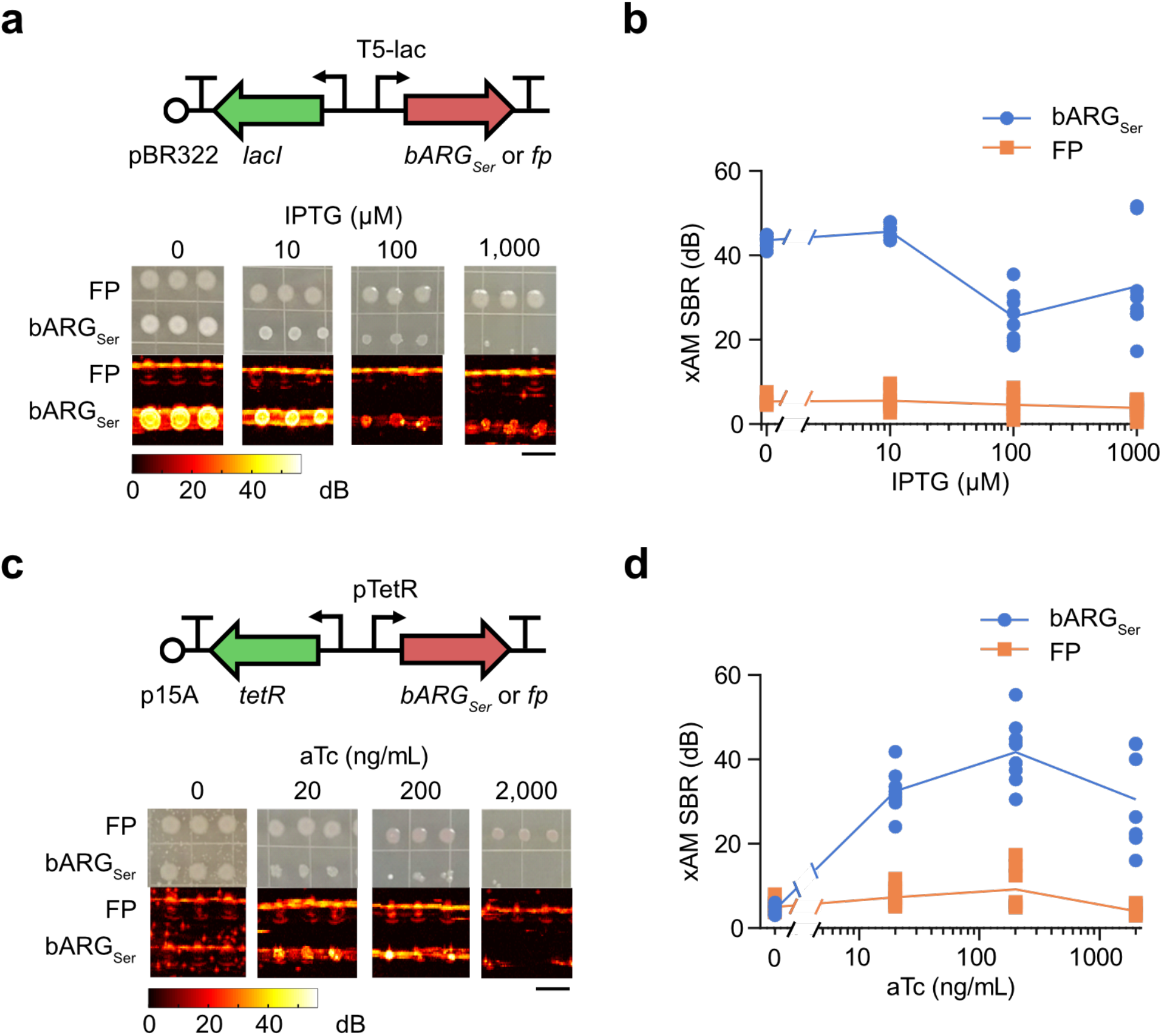
Testing bARG_Ser_ expression in EcN with IPTG- and aTc-inducible gene circuits. (**a**) Diagram of the IPTG-inducible construct used to express bARG_Ser_ in EcN (top), and representative optical and xAM images of bARG_Ser_-expressing or fluorescent protein (FP)-expressing patches of EcN on solid media with varying IPTG concentrations at 37°C (bottom). (**b**) Quantification of the xAM SBR of all patches from the experiment in (a) (n=8; curves represent the mean). (**c-d**) Same as in (a-b) but for the aTc-inducible construct. The scale bars in (a,c) represent 1 cm.

**Figure S10:**
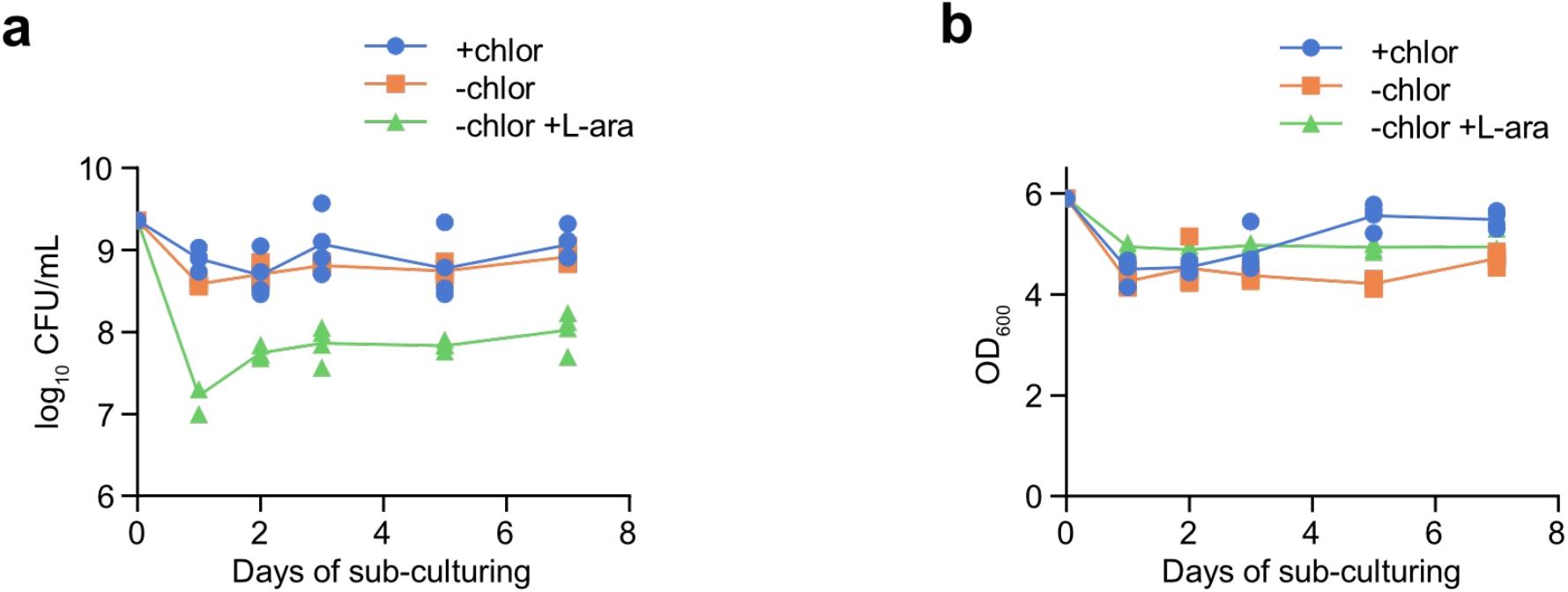
Effect of induction on viability and OD_600_ for bARG_Ser_-expressing EcN in liquid culture. (**a-b**) Colony forming units (CFU) per mL of culture (a) and optical density at 600 nm (b) during daily sub-culturing into LB media with 25 μg/mL chloramphenicol (+chlor), without chloramphenicol (chlor), or without chloramphenicol and with 0.1% (w/v) L-arabinose (−chlor +L-ara) using pBAD-bARG_Ser_-AxeTxe EcN. Curves represent the mean of n=4 biological replicates.

**Figure S11:**
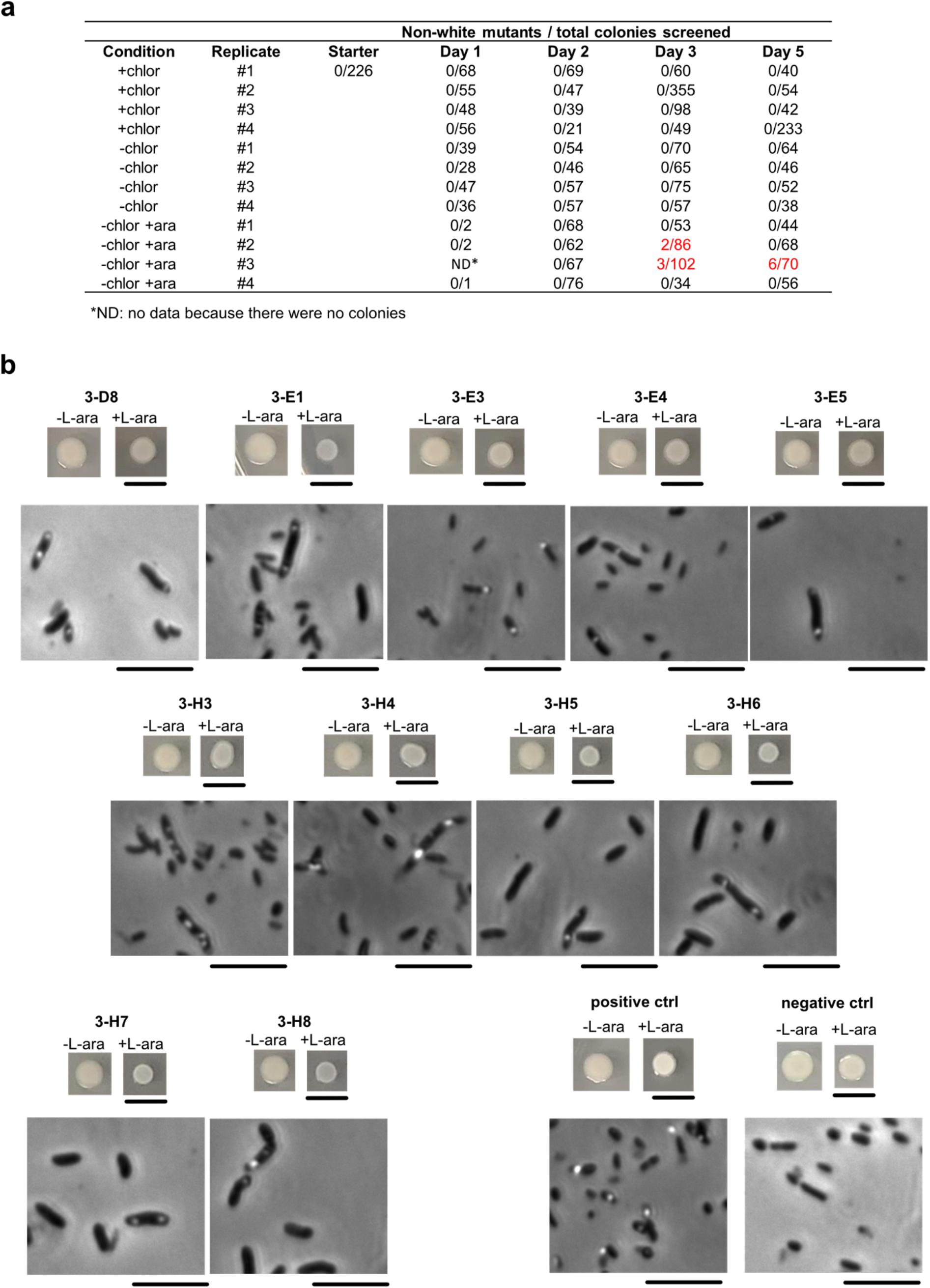
Quantification and characterization of EcN mutants deficient in bARG_Ser_ expression isolated from daily subculturing *in vitro*. (**a**) Numbers of non-white mutant colonies and total colonies screened on plates with 0.1% (w/v) L-arabinose from daily sub-culturing into LB media with 25 μg/mL chloramphenicol (+chlor), without chloramphenicol (−chlor), or without chloramphenicol and with 0.1% (w/v) L-arabinose (−chlor +L-ara) using pBAD-bARG_Ser_-AxeTxe EcN. Cultures where mutants were found are indicated in red. (**b**) Optical images of patches (top rows) on fresh plates with 0.1% (w/v) L-arabinose (+L-ara) and without L-arabinose (−L-ara), and phase contrast microscopy images (bottom rows) from the 11 mutant colonies in (a). Mutants 3-D3 and 3-E1 were from the culture −chlor +ara, replicate #2, day 3; mutants 3-E3, 3-E4, and 3-E5 were from the culture −chlor +ara, replicate #3, day 3; and mutants 3-H3 through 3-H8 were from the culture −chlor +ara, replicate #3, day 5. The positive and negative controls were wild-type pBAD-bARG_Ser_-AxeTxe EcN and pBAD-FP-AxeTxe EcN, respectively. Scale bars are 1 cm for images of patches and 10 μm for microscopy images.

**Figure S12:**
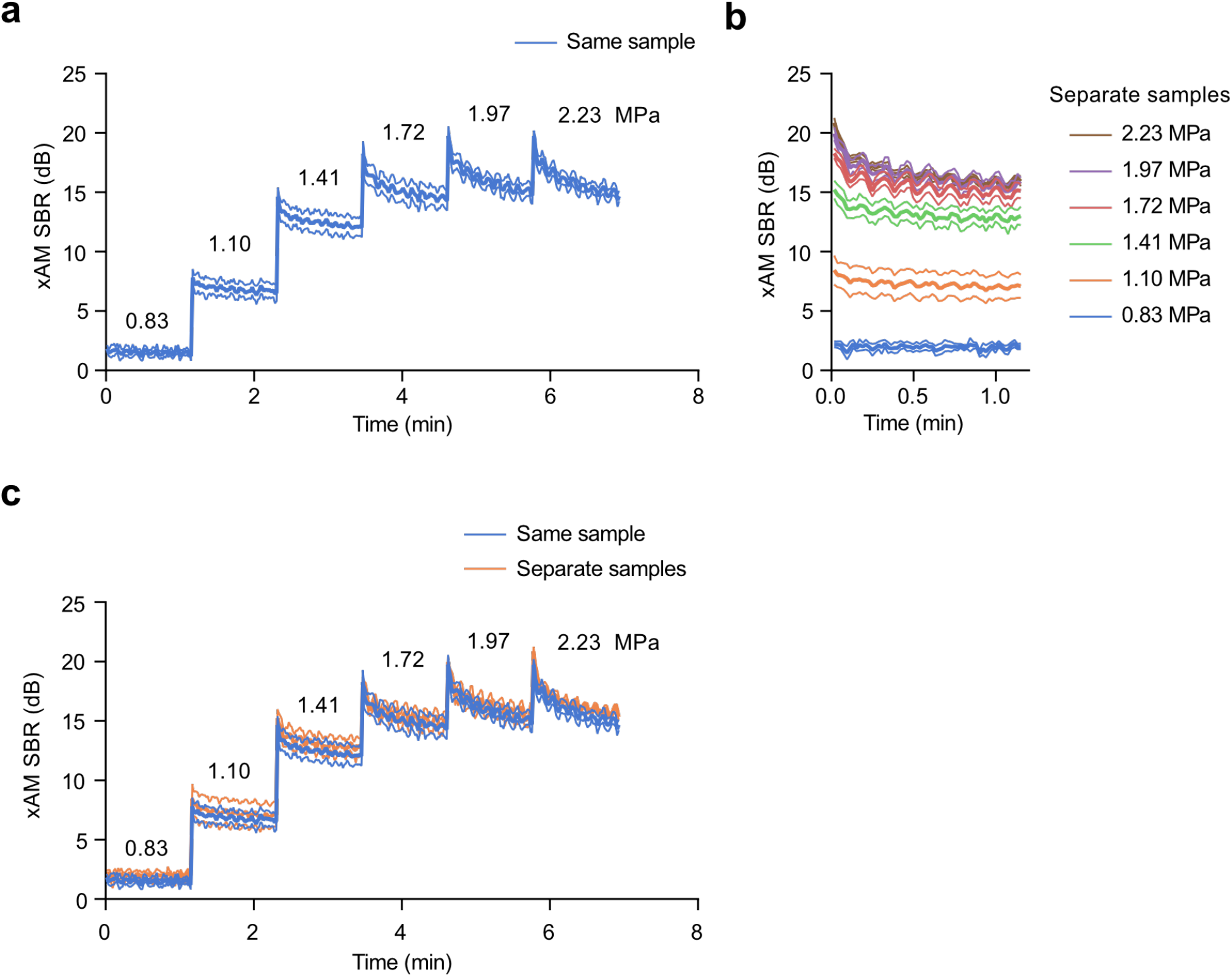
xAM ultrasound signal versus time at varying acoustic pressures applied sequentially to the same sample versus separate samples. (**a-b**) xAM SBR of bARG_Ser_-expressing EcN measured over time at various acoustic pressures. In (a), samples were subjected sequentially to 6 increasing acoustic pressures for approximately 70 sec each, whereas in (b) separate samples subjected to only one acoustic pressure for approximately 70 sec. (**c**) Overlay of xAM SBR curves for separate samples from (b) onto the curves for samples subjected to all pressures from (a). There is no difference between these curves, indicating that the xAM SBR measured at a certain pressure was not significantly affected by collapse at a previously applied pressure. For (a-c), pBAD-bARG_Ser_-AxeTxe EcN were induced with 0.1% (w/v) L-arabinose for 24 hours at 37°C in liquid culture, and were then normalized to 10^9^ cells/mL in agarose phantoms for ultrasound imaging. Bold lines represent the mean and thin lines represent ± standard deviation; n=3 biological replicates, each with 2 technical replicates. Imaging was performed with an L22-v14X transducer, so the values for pressure and xAM SBR do not exactly match those in Fig. 3a-c where an L22-v14 transducer was used.

**Figure S13:**
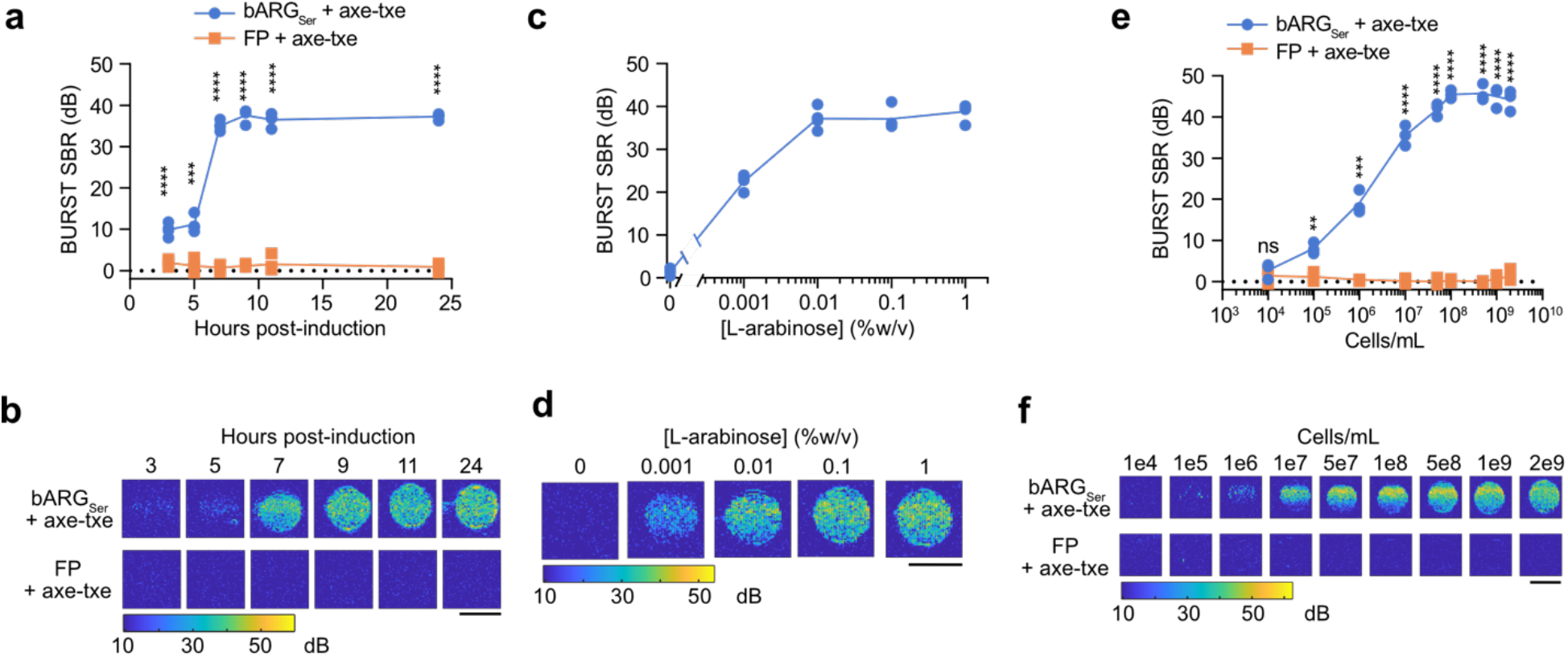
*In vitro* characterizations of bARG_Ser_-expressing EcN with BURST ultrasound imaging. (**a-b**) BURST ultrasound signal-to-background ratio (SBR) versus time after inducing pBAD-bARG_Ser_-AxeTxe and pBAD-FP-AxeTxe EcN strains with 0.1% L-arabinose in liquid culture at 37°C (a) and the corresponding representative BURST images (b). (**c-d**) BURST ultrasound SBR versus L-arabinose concentration used to induce pBAD-bARG_Ser_-AxeTxe EcN in liquid culture at 37°C for 24 hours (c) and the corresponding representative BURST images. (**e-f**) BURST ultrasound SBR versus concentration of pBAD-bARG_Ser_-AxeTxe or pBAD-FP-AxeTxe EcN cells induced for 24 hours at 37°C with 0.1% L-arabinose in liquid culture (e) and the corresponding representative BURST images (f). Note that the BURST SBR saturated at 7 hours post-induction, 0.01% (w/v) L-arabinose, and 10^8^ cells/mL. All scale bars are 2 mm. For (a-d), cells were normalized to 10^9^ cells/mL in agarose phantoms for ultrasound imaging. For (a, c, e), each point is a biological replicate (n=4 for a and c; n=3 for e) that is the average of at least 2 technical replicates and curves indicate the mean. Asterisks represent statistical significance by two-tailed, unpaired Student’s t-tests (**** = p<0.0001, *** = p<0.001, ** = p<0.01, ns = no significance).

**Figure S14:**
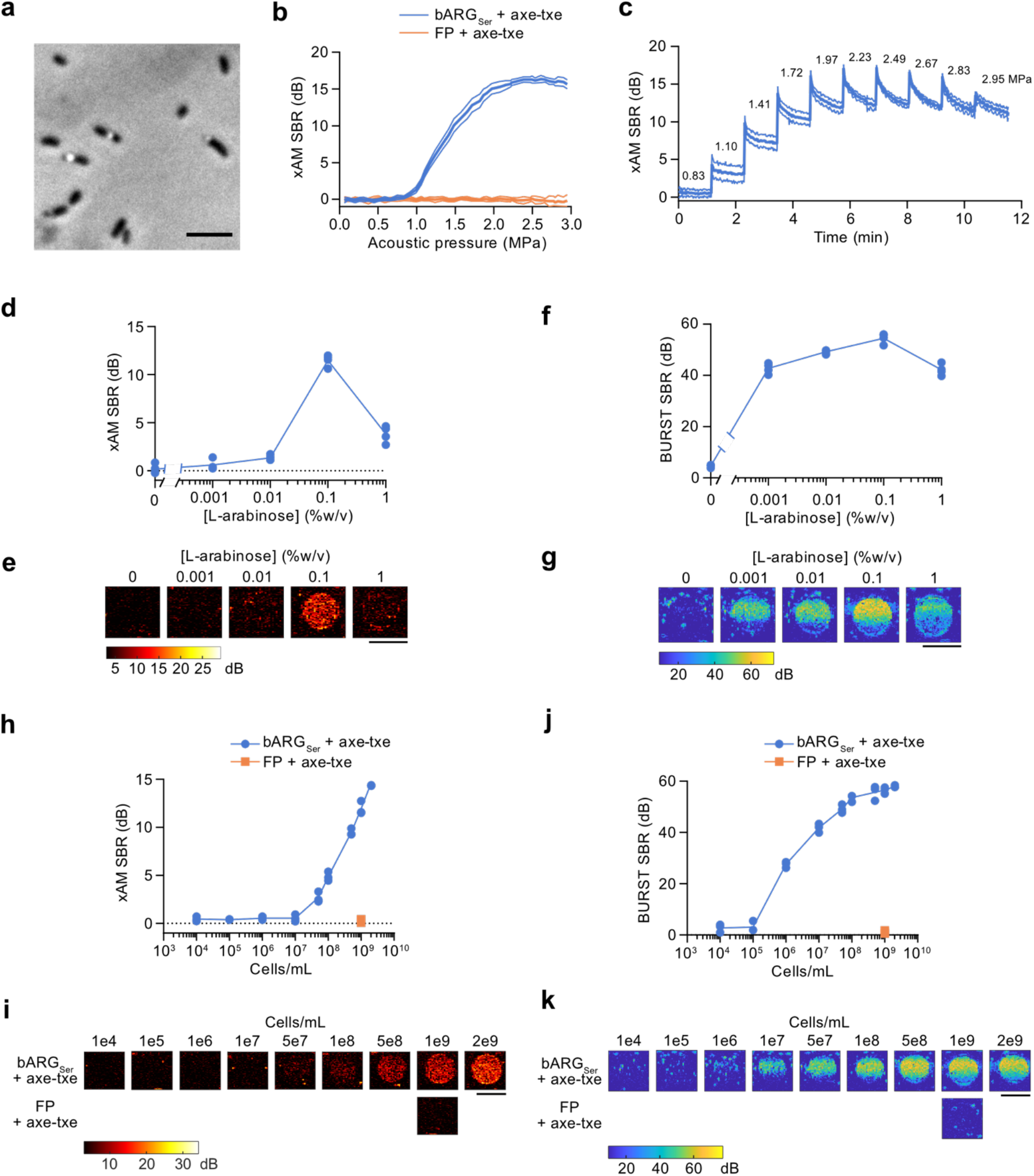
bARG_Ser_ expression and acoustic characterization in *Salmonella enterica* serovar Typhimurium. (**a**) Representative phase contrast microscopy image of bARG_Ser_-expressing *S. Typhimurium* cells. (**b**) xAM SBR as a function of transmitted acoustic pressure for bARG_Ser_-expressing and FP-expressing *S. Typhimurium* cells. (**c**) xAM SBRs measured over time when the transmitted acoustic pressure was increased approximately every 70 sec as indicated by the numbers above the curve for bARG_Ser_-expressing *S. Typhimurium.* Ultrasound was applied at a pulse repetition rate of 86.8 Hz. For (a-c), pBAD-bARG_Ser_-AxeTxe *S. Typhimurium* cells were induced with 0.1% (w/v) L-arabinose for 24 hours at 37°C in liquid culture, and were then diluted to 10^9^ cells/mL in agarose phantoms for ultrasound imaging. Bold lines represent the mean and thin lines represent ± standard deviation; n=3 biological replicates, each with 2 technical replicates. (**d-g**) xAM ultrasound SBR (d) and corresponding representative ultrasound images (e), and BURST SBR (f) and corresponding representative images (g), using varying L-arabinose concentrations to induce pBAD-bARG_Ser_-AxeTxe *S. Typhimurium* in liquid culture at 37°C for 24 hours. (**h-k**) xAM ultrasound SBR (h) and corresponding representative ultrasound images (i), and BURST SBR (j) and corresponding representative images (k), of varying concentrations of pBAD-bARGs_er_-AxeTxe or pBAD-FP-AxeTxe *S. Typhimurium* cells induced for 24 hours at 37°C with 0.1% (w/v) L-arabinose in liquid culture. Scale bars represent 5 μm in (a) and 2 mm in (e, g, i, k). For (d-g), cells were diluted to 10^9^ cells/mL in agarose phantoms for ultrasound imaging. For (d, f, h, j), each point is a biological replicate (n=4 for d and f; n=3 for h and j) that is the average of at least 2 technical replicates, and curves represent the mean. All ultrasound imaging in this figure was performed with an L22-14vX transducer.

**Figure S15:**
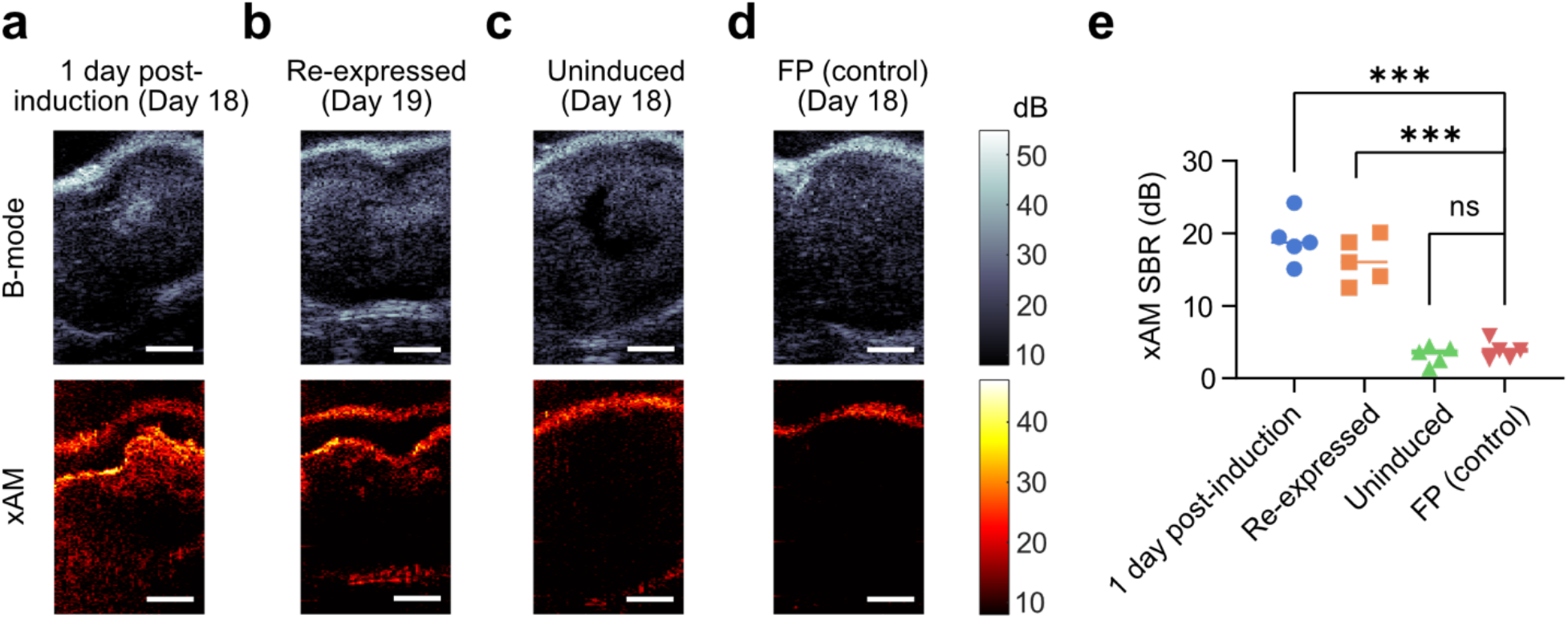
xAM ultrasound imaging of mouse tumors colonized by EcN. (**a-c**) Representative B-mode (top, grayscale) and xAM (bottom, hot-scale) ultrasound images of tumors colonized by pBAD-bARGs_er_-AxeTxe EcN at least 24 hours after induction with L-arabinose on day 18 (a), at least 24 hours after collapse and re-induction (day 19) (b), or uninduced on day 18 (c). (**d**) Representative ultrasound images of tumors colonized by pBAD-FP-AxeTxe EcN at least 24 hours after induction with L-arabinose on day 18. Scale bars in (a-d) represent 2 mm. (**e**) Quantification of the xAM SBR for the same conditions in (a-d). Each group is n=5 mice and lines indicate the mean. Asterisks represent statistical significance by two-tailed, unpaired Student’s t-tests (*** = p<0.001, ns = no significance). See **Fig. 4a** for the corresponding *in vivo* protocol.

**Figure S16:**
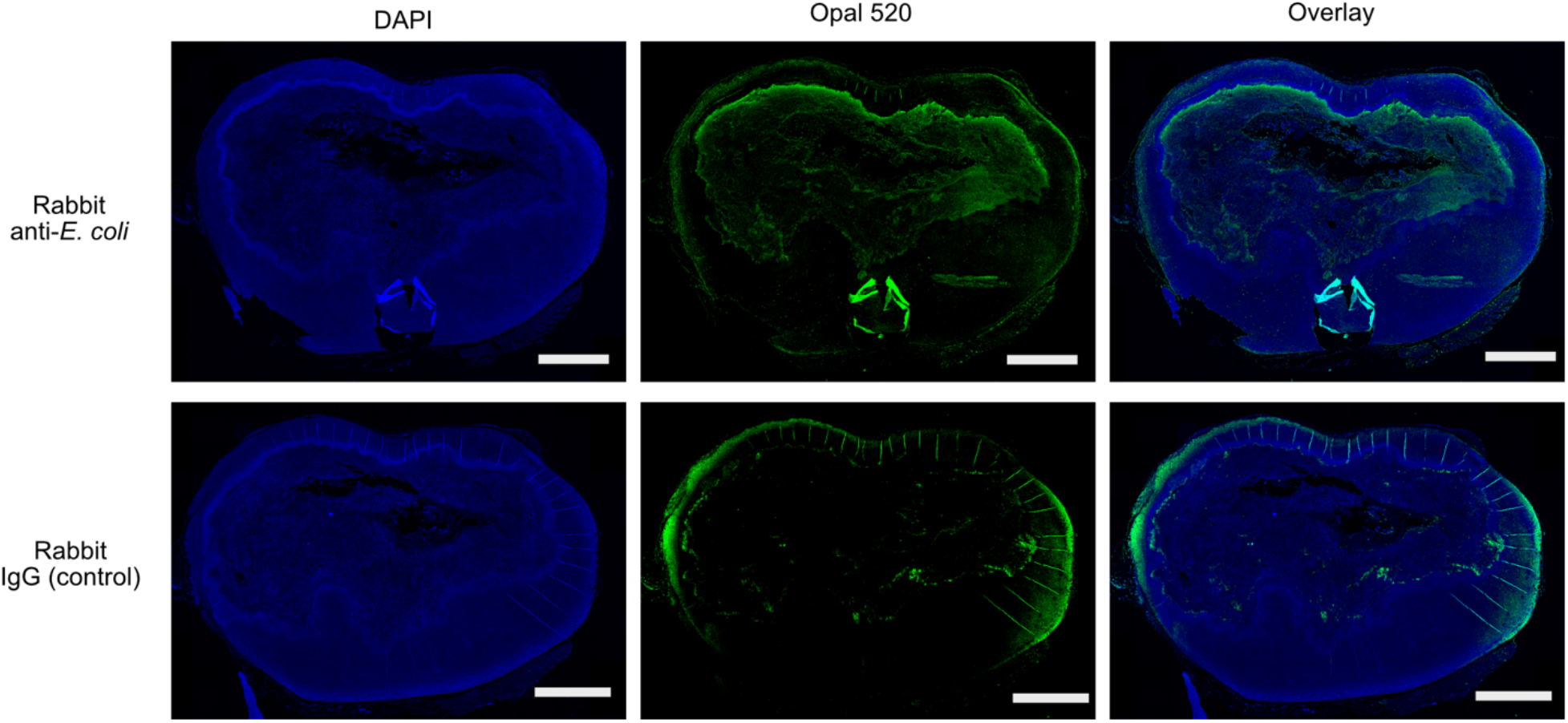
Histology of MC26 tumor colonized with bARG_Ser_-expressing EcN. Fluorescent images of tissue sections after ultrasound imaging on day 19 (see **Fig. 4a**). Sections were incubated with either polyclonal rabbit anti-*E. coli* antibodies (top row) or non-reactive rabbit IgG isotype control antibody (bottom row) as a negative control. All sections were then incubated with an Opal 520 polymer anti-rabbit HRP antibody (Akoya biosciences) and counterstained with DAPI. The EcN are visible in the necrotic core in the Opal 520 channel (top middle panel); the edges of the tissue exhibit a high degree of background staining (bottom middle panel).

**Figure S17:**
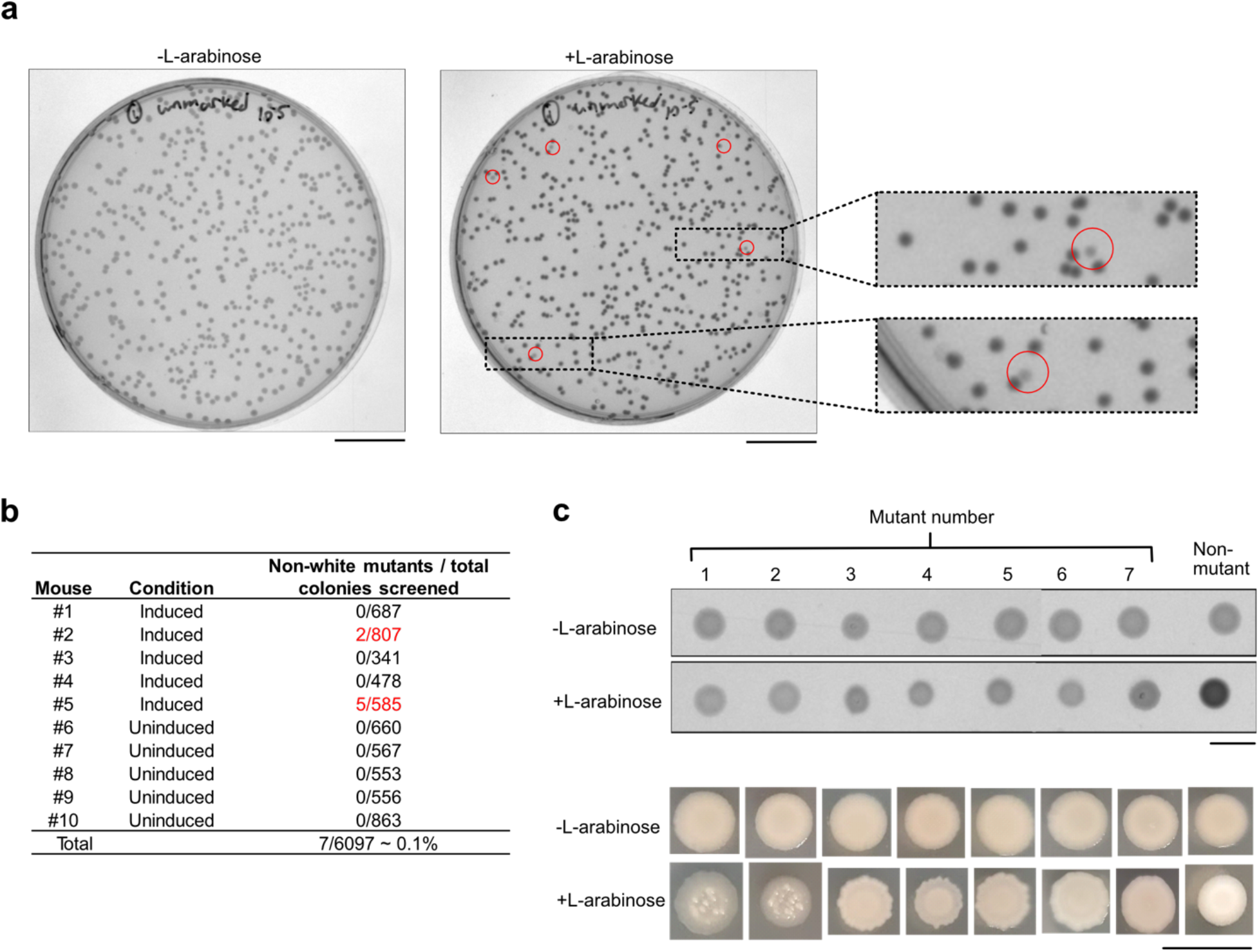
Screening for EcN mutants defective in bARG_Ser_ expression isolated from colonized tumors. (**a**) White light transmission images of plates with 0.1% (w/v) L-arabinose and without L-arabinose from plating a tumor (from mouse #5 in (b)) colonized by bARG_ser_-expressing EcN. Mutant colonies on the +L-arabinose plate appear lighter (more translucent) than wild-type opaque colonies and are indicated by red circles. (**b**) Numbers of non-white mutant colonies and total colonies screened on plates with 0.1% (w/v) L-arabinose for the ten mice injected with pBAD-bARG_ser_-AxeTxe EcN. **c**, White light transmission images (top) and photographs (bottom) of patches on fresh plates with 0.1% (w/v) L-arabinose and without L-arabinose made from the seven translucent mutant colonies in red in (b) and an opaque nonmutant colony as a control. Mutants 1-2 were from mouse #2 and mutants 3-7 were from mouse #5. Scale bars are 2 cm in (a) and 1 cm in (c).

**Figure S18:**
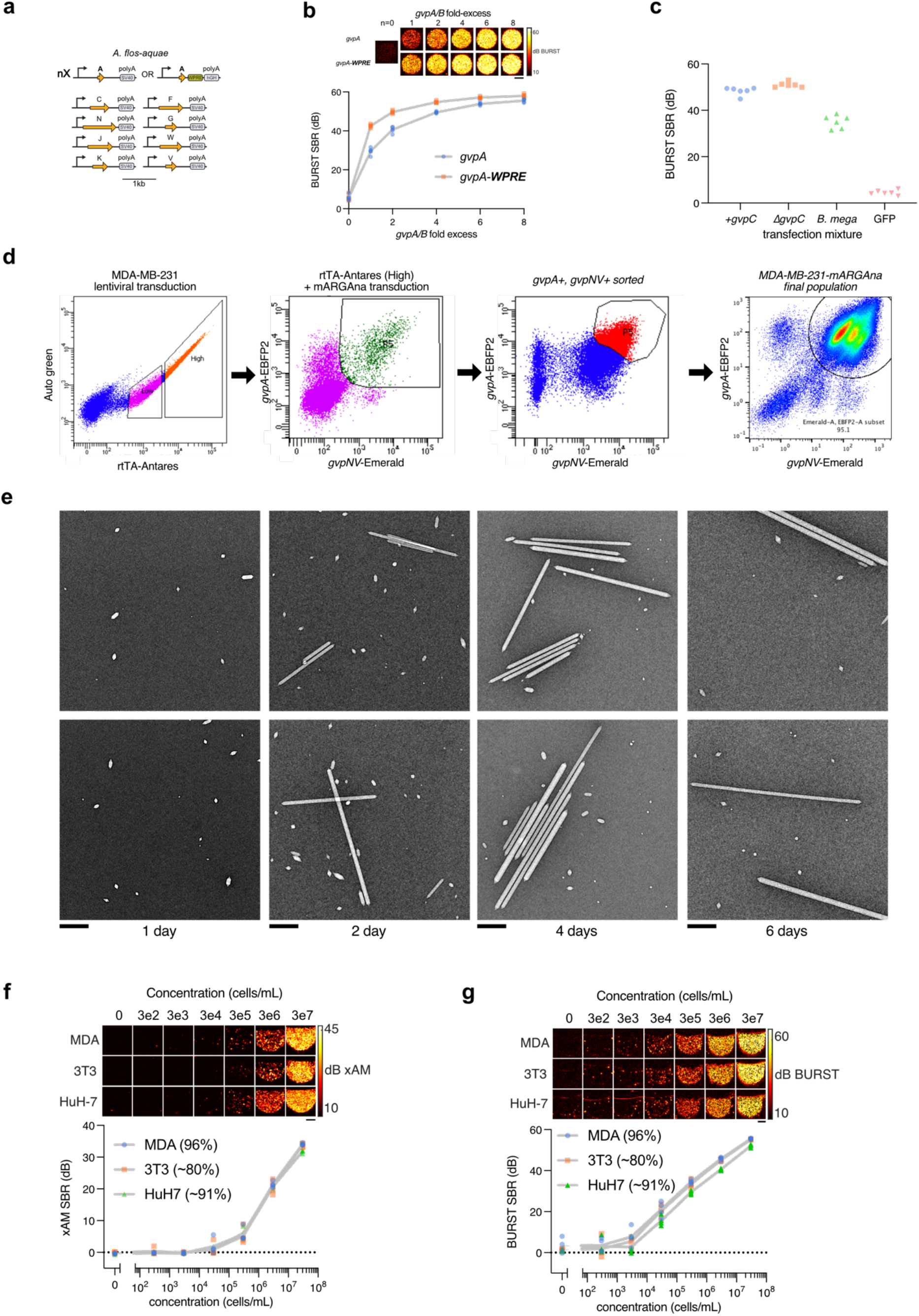
Additional data on heterologous expression of the *A. flos-aquae* GV gene cluster in mammalian cells. (**a**) Schematic of the codon-optimized *A. flos-aquae* monocistronic plasmid set used in this study. (**b**) Representative BURST images (top) and SBR quantification (n=5, bottom) of transient GV expression in HEK293T cells 3 days after co-transfection of mixtures with varying *gvpA* fold excess relative to their respective assembly factor plasmids, with and without WPRE elements on the *gvpA* DNA. Gray lines connect the means of replicates. (**c**) BURST SBR quantification (n=6) of 293T cells expressing constructs tested in **Fig. 5d**. (**d**) FACS and flow cytometry data for production of the MDA-MB-231-mARG_Ana_ cell line. rtTA expressing “High” cells were collected for subsequent mARG/transposase transduction. Cells expressing *gvpA* and *gvpNV* were sorted twice to arrive at the final ~95% pure population. (**e**) TEM images of GVs purified from MDA-MB-231-mARG_Ana_ detergent lysates after 1, 2, 4 and 6 days of 1 μg/ml doxycycline induction. Scalebars represent 0.5 μm. (f) Representative xAM images (top) and SBR quantification (n=4, bottom) of induced MDA-MB-231-mARG_Ana_, 3T3-mARG_Ana_ and HuH7-mARG_Ana_ cells at 0.61 MPa as a function of cell concentration. Limit of detection was 300k cells/mL for MDA-MB-231 and 3T3, and 30k cells/mL for HuH7 with p<0.05 by unpaired t-tests. (g) Representative BURST images (top) and SBR quantification (n=4, bottom) of induced MDA-MB-231-mARG_Ana_, 3T3-mARG_Ana_ and HuH7-mARG_Ana_ cells as a function of cell concentration. Limit of detection was 30k cells/mL for MDA-MB-231 and HuH7, and 3k cells/mL for 3T3 with p<0.05 by unpaired t-tests. For (f) and (g), gray lines connect the means of the replicates and scalebars represent 1 mm. Percentages in parentheses represent mARG_Ana_-positive cells.

**Figure S19:**
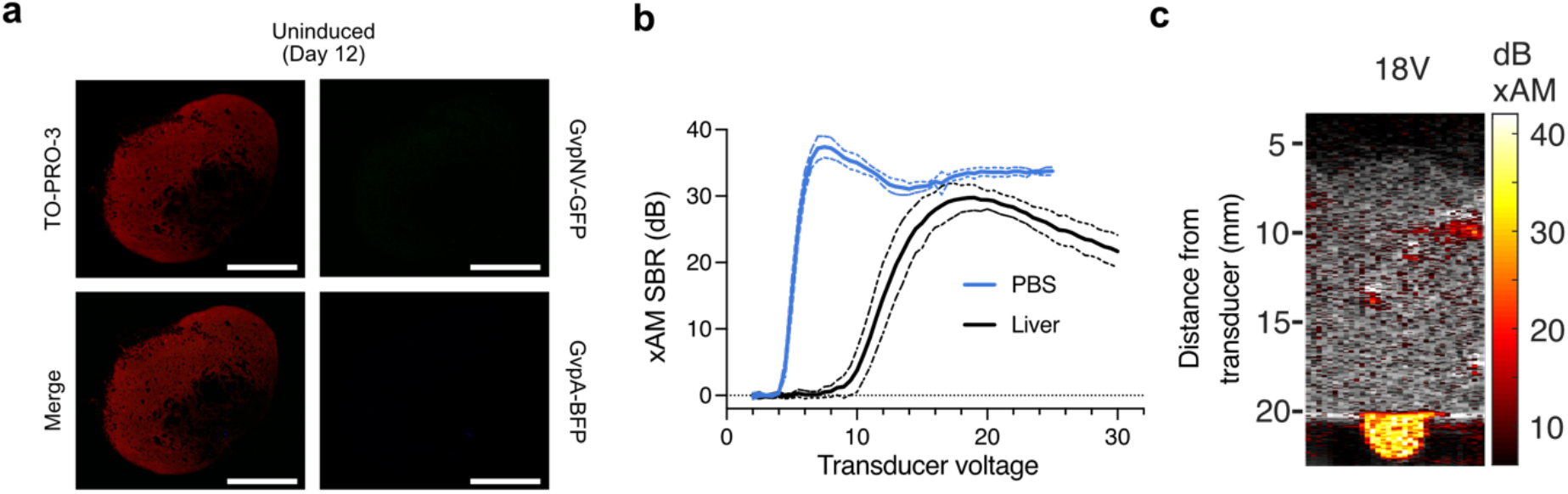
Tumor histology and imaging through thick tissue. (**a**) Fluorescence micrograph of a 100 nm-thick tissue section from an uninduced tumor. The red color shows TO-PRO-3 nuclear stain; GFP and BFP fluorescence were not observed for this control tumor. Scale bars represent 2 mm. (**b**) Quantification of xAM SBR of MDA-MB-231-mARG_Ana_ cells imaged under >1 cm of beef liver tissue (n=7) or under PBS (n=4) as a function of transducer voltage. Thick lines represent the mean of replicates and thin lines represent ± standard deviation. (**c**) Representative xAM/B-mode overlay of MDA-MB-231-mARG_Ana_ cells imaged under liver tissue using L10-5v transducer at 18V for xAM.

**Figure S20:**
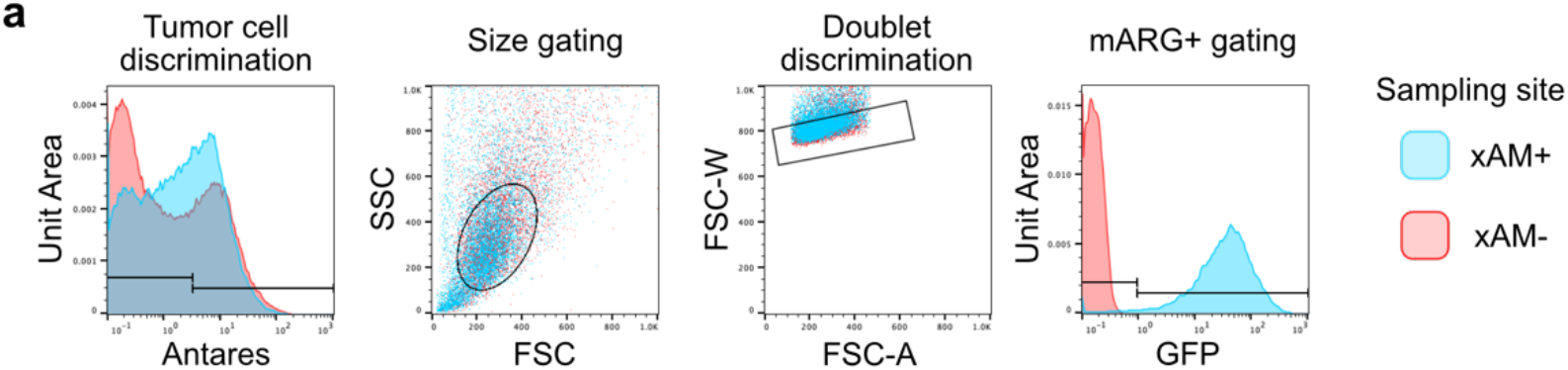
Flow cytometric gating strategy for chimeric tumor biopsy sample analyses. Events were first gated based on Antares expression to exclude endogenous mouse cells. Antares-positive cells were then gated by size. Single cells were gated based on FSC-W vs FSC-A plot. GFP-positive (mARG_Ana_-positive) cells were gated from the resulting histograms.

**Supplementary Video 1: xAM/B-mode tomogram of an induced orthotopic tumor imaged on day 12.** A representative tomogram of an orthotopic MDA-MB-231-mARG_Ana_ tumor imaged after 12 days of doxycycline induction. Each slice in the tomogram is separated by 100 μm.

**Supplementary Video 2: xAM/B-mode tomogram of an induced chimeric tumor imaged on day 5.** A representative tomogram of a chimeric MDA-MB-231-mARG_Ana_ tumor imaged after 5 days of doxycycline induction. Each slice in the tomogram is separated by 200 μm.

**Supplementary Video 3: xAM/B-mode 3D reconstruction of an induced chimeric tumor imaged on day 5.** 3D B-mode and xAM data were smoothened and converted to isosurfaces using Matlab. Yellow 3D map represents B-mode density; blue 3D map represents xAM density.

**Supplementary Video 4: Representative xAM/B-mode video of a chimeric tumor biopsy procedure sampling the xAM-positive region.**

**Supplementary Video 5: Representative xAM/B-mode video of a chimeric tumor biopsy procedure sampling the xAM-negative region.**

## Notes

https://tinyurl.com/arg2vids

### Competing Interest Statement

The authors have declared no competing interest.

### Summary of Updates

This revision contains additional demonstrations supporting the broad utility of second-generation acoustic reporter genes (ARGs). - Demonstrating the use of second-generation ARGs in real-time ultrasound-guided biopsies. - Demonstrating the imaging of mammalian ARGs through deeper tissues. - Demonstrating the use of mammalian ARGs in additional cell types (HuH7 and NIH-3T3). - Demonstrating the use of bacterial ARGs in additional species (S. typhimurium).

